# Long-range allosteric restraint by the Spiral–Pincer scaffold gates oxidative activation of the ATM kinase

**DOI:** 10.64898/2026.11.02.763569

**Authors:** Anirudh Chandra, Biswajit Sadhu

## Abstract

ATM is a master regulator of the DNA-damage response, activated either by DNA double-strand breaks through the MRN complex or, independently, by oxidative stress, which drives disulfide bond formation between Cys2991 residues on the two halves of the ATM dimer. Although both pathways involve release of the PIKK regulatory domain (PRD) from the kinase active site, how the distant Spiral–Pincer scaffold regulates access to this state remains unclear, partly because the PRD–FAT-TRD3 interface is unresolved in available experimental structures. Here, we combine an AlphaFold3-completed full-length ATM dimer model with well-tempered metadynamics to map its conformational landscape. We find that an oxidation-competent conformation already exists in the resting dimer, although it is rarely populated, supporting conformational selection rather than induction of a new state. Removing Spiral–Pincer eliminates this restriction: the conformation becomes dominant, and the landscape narrows to a few well-separated states. The dimer’s two halves respond unequally, one loop of the PRD rearranging substantially while the other remains unchanged, via specific domain contacts rather than a direct route to the reactive cysteines. This identifies Spiral–Pincer as a release-controlled brake on ATM activation, a mechanism that may extend across the wider PIKK kinase family.

## Introduction

Ataxia-telangiectasia mutated (ATM) is a serine/threonine kinase of the phosphatidylinositol-3-kinase-like kinase (PIKK) family that coordinates the cellular response to genomic stress by phosphorylating downstream effectors controlling DNA repair, cell-cycle arrest, and apoptosis [1–4]. Loss-of-function mutations in ATM cause the genome-instability disorder ataxia-telangiectasia [5]. This activation of the ATM homodimer happens through at least two structurally distinct routes. Following DNA double-strand breaks, the Nbs1 of the Mre11–Rad50–Nbs1 (MRN) complex engages the N-terminal region of ATM and promotes a rearrangement of the autoinhibited dimer that favours catalytically active monomers [6]. Oxidative stress, in contrast, activates ATM while it remains dimeric, through formation of an intermolecular disulfide bond between Cys2991 residues contributed by each protomer [7]. That two mechanistically unrelated stimuli converge on kinase activation indicates that ATM does not possess a single activation pathway; rather, its architecture must permit access to a shared catalytically competent state from multiple structurally different starting points.[7–9] The central question is therefore not simply how ATM becomes active, but how its overall architecture regulates *access* to the substrate-binding site, and which conformational states are available to each activating signal.

Structural studies have established that the ATM kinase domain retains a largely catalytically competent fold even in the autoinhibited state [8, 10–13]. Substrate access to this catalytic cleft is instead restricted by a regulatory assembly: the FRAP, ATM, TRRAP (FAT) domain, the PIKK regulatory domain (PRD), the FATC domain, and part of the kinase C-lobe. Together, these elements occlude the substrate-binding groove on the kinase activation loop [10, 13, 14] (Fig. 1). In the inactive ATM dimer, contacts between the FAT and Kinase domains of opposing protomers stabilize the PRD in an occluding position; relaxation of these contacts is accompanied by PRD disorder and increased accessibility to the catalytic cleft [8, 13, 14]. An important, unresolved component of ATM activation is therefore how this local rearrangement is regulated by the much larger architecture of the protein.

**Figure 1:**
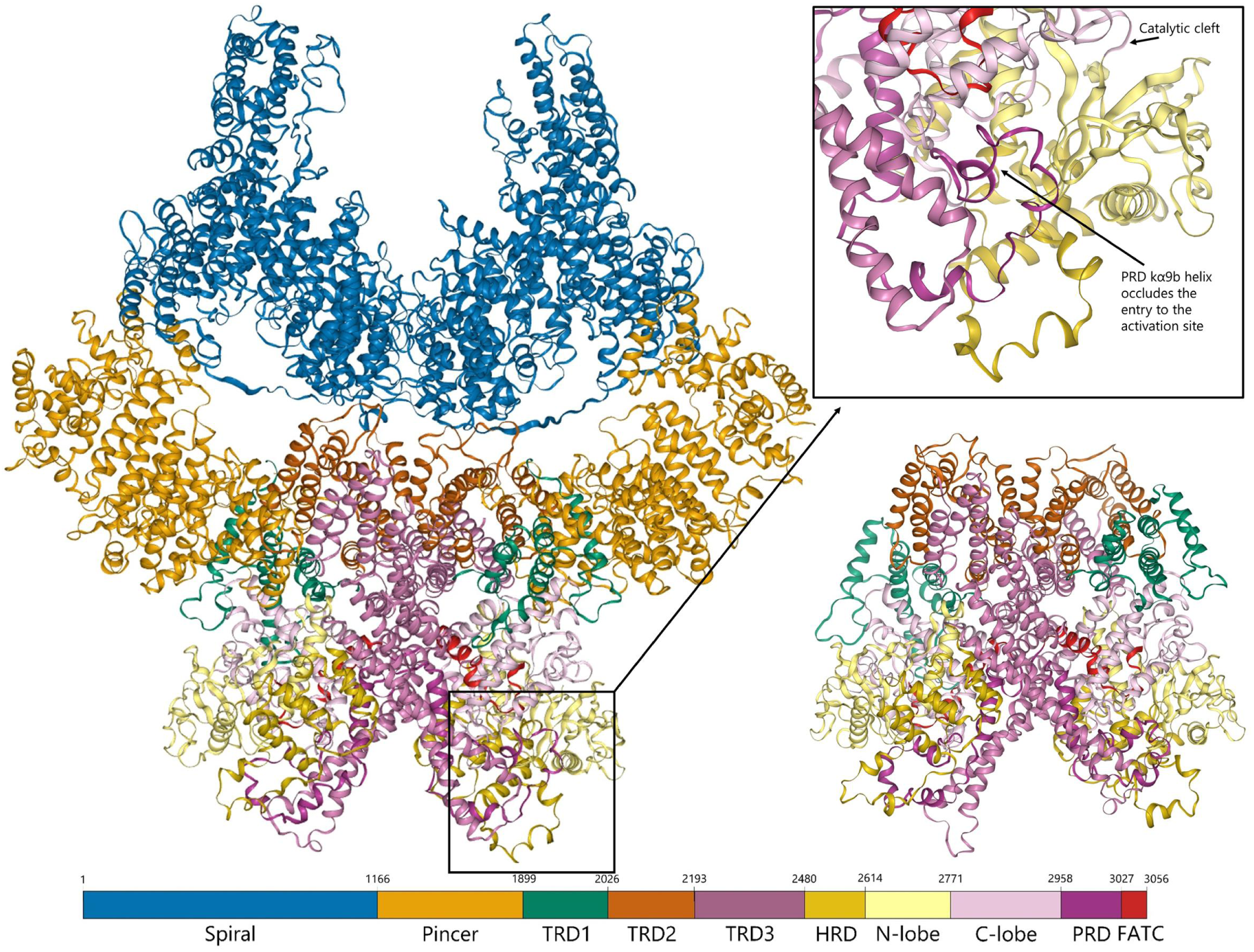
Domain architecture of the human ATM homodimer, full-length and Spiral–Pincer-truncated. Structure (left and right) and linear domain map with residue boundaries (bottom) of one ATM protomer within the dimer, for the full-length construct (left) and the Spiral–Pincer-truncated construct (right) used for the causal truncation experiments described below. Domain boundaries follow the convention established for the experimental ATM structure [13].

It is well-documented that the far-distant domains N-terminal Spiral and Pincer domains from the catalytic core, are extended *α*-solenoids, that form the principal interaction platform for MRN and other regulatory factors. They sit almost 100 Å from the kinase activation site [13–15], yet the Pincer packs directly against the FAT–kinase domains – a plausible mechanical route by which motion of the N-terminal scaffold could influence the catalytic domain. Structural studies report substantial Spiral/Pincer mobility [13, 15], and a recent cryo-EM study captured coordinated displacement of the Spiral, Pincer, and kinase N-lobe accompanying PRD and cleft reorganization during activation [16]. These observations do not, however, establish whether Spiral–Pincer motion is intrinsically coupled to active-site accessibility or reflects independent fluctuations captured incidentally in static snapshots – indeed, one earlier structure [13] showed substantial N-terminal mobility with little accompanying FAT–kinase rearrangement. This raises the central mechanistic question: does the Spiral–Pincer scaffold merely move, or does its motion actively reshape the energetic landscape governing access to substrate-compatible kinase conformations?

This question bears directly on oxidative activation, because Cys2991, the key residue behind the S-S covalent bond formation, lies within a flexible loop of the PRD, the same regulatory element that gates the substrate-accessible catalytic cleft. The PRD is proposed to occlude this cleft by packing against the FAT-TRD3 domain of the opposing protomer [8, 10], but this interaction has never been directly visualized: both the PRD and FAT-TRD3 have unresolved residues at exactly this interface in every deposited ATM structure [8, 10–13] (Supplementary Table S1). This leaves one fundamental question unanswered: is the oxidized, disulfide-linked conformation generated *de novo* by covalent bond formation, or does it instead represent a pre-existing state of the ATM dimer’s intrinsic conformational ensemble that oxidation merely stabilizes?

We hypothesized that long-range coupling between the Spiral–Pincer scaffold and the FAT– kinase domains produces a heterogeneous free-energy landscape, one containing metastable, partially accessible substrate-compatible states. On this view, the N-terminal scaffold itself regulates the population and accessibility of these states – functioning as an allosteric restraint on activation rather than as a conduit for an activating signal.

Testing this hypothesis requires resolving conformational transitions that are difficult to access experimentally and unlikely to be adequately sampled by conventional molecular dynamics. We therefore developed an atomistic model of the full-length human ATM dimer by integrating experimentally resolved regions with AlphaFold-based completion of unresolved regions, including the Cys2991-containing PRD loop, and characterized its conformational landscape using well-tempered metadynamics. We identify a pre-existing, metastable conformation in the native dimer that is compatible with Cys2991 disulfide formation and show, through Spiral–Pincer truncation (Fig. 1), that the distal scaffold controls its energetic accessibility. Comparison with experimentally determined inactive and oxidized structures, together with domain- and residue-resolved interaction-network analysis, further reveals that access to this state is accompanied by asymmetric PRD remodelling and long-range reorganization of the FAT–kinase–PRD architecture rather than a direct residue-level connection to Cys2991. These results support a restraint-release mechanism in which the Spiral–Pincer scaffold limits access to a pre-existing oxidation-compatible conformational ensemble.

## Results

### An AlphaFold3-completed model reconstructs the previously unresolved PRD–FAT-TRD3 interface implicated in substrate occlusion

A major challenge in investigating the molecular mechanisms of ATM, a 6,112-residue homodimer, is the lack of a complete, well-resolved experimental structure. In particular, the mechanistically important PRD and FAT-TRD3 domains are not simultaneously resolved at their proposed mutual interface in available deposited ATM structures, precluding direct visualization of the proposed substrate-occlusion contact [8, 10]. To address this limitation, we generated an atomistic model of the full-length human ATM dimer using AlphaFold3. Five structural candidates were screened for steric clashes and predicted disorder, and the candidate showing the closest global and domain-wise agreement with the experimental structure was selected (Supplementary Fig. S1; Supplementary Tables S4–S6). The selected model reconstructs the PRD and FAT-TRD3 simultaneously, providing a structural framework in which their proposed substrate-occlusion interface can be examined (Supplementary Fig. S6). Support for the structural plausibility of this region comes from its closer domain-wise agreement with the experimental structure relative to other regions of the model (Supplementary Table S5). Notably, FAT-TRD3 exhibits reasonably high per-residue confidence scores (pLDDT), whereas the PRD has comparatively lower confidence, consistent with its proposed intrinsic disorder [17, 18] and its poor resolution in the experimental structure. Further details of AlphaFold3 model generation, candidate selection, and structural validation are provided in Supplementary Note 1 and Supplementary Table S7.

### An oxidation-compatible conformation pre-exists within the native free-energy land-scape of full-length ATM

The signature of oxidised ATM activation is the formation of cross-protomer Cys2991–Cys2991 disulfide bond. To determine whether the geometry required for the disulfide bond formation is intrinsic to the native conformational ensemble of ATM or is instead generated only upon covalent bond formation, we reconstructed the two-dimensional free-energy surface (FES) of the full-length ATM dimer using well-tempered metadynamics. The FES was reconstructed along two collective variables: the inter-protomer separation of the Cys2991 residues (CV1) and the relative positioning of the PRD with respect to the substrate-binding site of the kinase activation loop (CV2).

The resulting landscape is rugged (Fig. 2a), containing multiple well-defined local minima separated by comparatively shallow barriers. Watershed segmentation identified eight major metastable basins (F1–F8). The experimentally determined inactive ATM structure (PDB 7SIC) [13] falls within the global free-energy basin, in close proximity to the three lowest-energy minima (F1–F3), which differ from one another by less than 0.3 kcal mol*^−^*^1^ and are therefore expected to interconvert freely at physiological temperature. Therefore, the inactive crystallographic/cryo-EM structure possibly represents one member of a broad, thermally accessible ensemble rather than a single discrete conformation.

**Figure 2:**
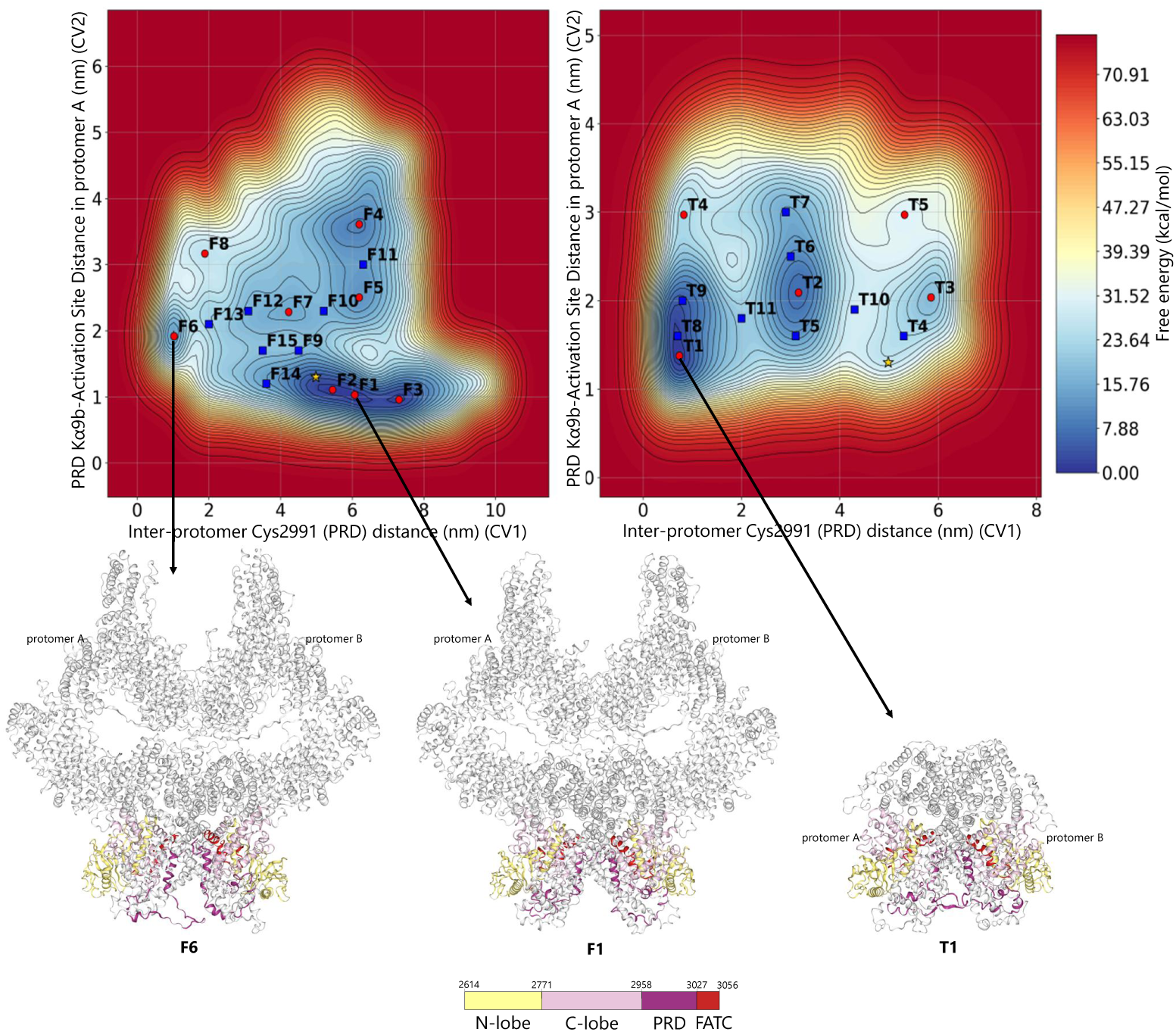
The free-energy landscape of full-length ATM contains multiple metastable states, including an oxidation-compatible basin (F6). Two-dimensional free-energy surface of (a) the full-length (top left) and (b) the Spiral–Pincer-truncated ATM dimer (top right), along CV1 (inter-protomer Cys2991–Cys2991 distance) and CV2 (PRD–activation-loop distance), reconstructed by well-tempered metadynamics. Red circles are local minima (F1–F8 in (a); T1–T5 in (b)) identified by watershed segmentation algorithm; blue squares are local maxima/saddle points; yellow star is the position of the experimentally determined inactive ATM structure (PDB 7SIC) [13].

Beyond this native ensemble, the FES reveals several previously uncharacterized metastable states. F4, the deepest and most extensively sampled of these, lies 8.5 kcal mol*^−^*^1^ above the global minimum. Most notably, a separate basin, F6, closely reproduces the relative PRD arrangement of the experimentally determined oxidized ATM structure (PDB 8OXM) [8] – quantified in a later section (Supplementary Table S8) – bringing the two Cys2991 residues into a proximity compatible with intermolecular disulfide bond formation. F6 lies 12.8 kcal mol*^−^*^1^ above the global minimum and is therefore only transiently populated under resting conditions (convergence of this and all other reported basin free energies is assessed in Methods; see Supplementary Fig. S8). Recovering this conformation without imposing a disulfide bond or any oxidation-specific restraint indicates that the oxidation-compatible geometry is encoded within the intrinsic dynamics of the inactive dimer, and that oxidative stress most plausibly acts by stabilizing (via covalent cross-linking) an otherwise rare, pre-existing state – a conformational-selection mechanism, rather than one in which disulfide formation induces a conformation absent from the resting ensemble.

### The Spiral–Pincer scaffold restrains access to the oxidation-compatible state

The FES shows that the oxidation-compatible geometry is rare but real – a state the native dimer visits only reluctantly. We next asked what keeps it rare. The Spiral and Pincer *α*-solenoids sit more than 100 Å from Cys2991, yet prior work has proposed them as the platform through which distal regulatory inputs reach the kinase domain [13, 14]. If Spiral–Pincer actively transmits an activating signal, as expected for a conventional allosteric pathway, removing it should make the oxidation-compatible state *harder* to reach or leave it unchanged; if instead it passively restrains access to a state the kinase domain can already adopt on its own, removing it should make that state *easier* to reach. We tested this directly by reconstructing the free-energy landscape of a Spiral–Pincer-truncated ATM model under identical metadynamics conditions (Fig. 2b).

Removing Spiral–Pincer fundamentally reorganizes the conformational landscape. The near-degenerate F1–F3 basin, corresponding to the inactive state, is absent in the truncated protein: the global minimum (T1) instead corresponds to a compact, oxidation-compatible-like geometry, and every remaining minimum is now well separated, sitting at least 6.1 kcal mol*^−^*^1^ from its neighbours. The experimentally determined inactive ATM structure, which coincides with the dominant lowenergy basin in the full-length landscape, instead falls in a high-energy region of the truncated one. Deleting Spiral–Pincer, in other words, does not merely remove a correlated conformational change – it eliminates the 12.8 kcal mol*^−^*^1^ energetic penalty that keeps the oxidation-compatible geometry rare in the intact protein. This is potentially the signature of *restraint release* in which the Spiral–Pincer scaffold functions akin to an active energetic brake on PRD geometry.

### Distinct metastable ensembles are distinguished by reorganization of the regulatory scaffold, not the catalytic core

If Spiral–Pincer indeed acts as an energetic brake, its removal should not just change the thermodynamic populations of states – it should reveal *where* the released conformational freedom is expressed structurally. We therefore asked whether the metastable substates identified in the FES primarily involve the kinase core or instead arise from rearrangement of the surrounding regulatory architecture, including the PRD, FATC, and FAT domains, particularly FAT-TRD3. Previous structural studies have similarly shown that the ATM kinase core can remain well ordered while the surrounding FAT–PRD regulatory architecture undergoes substantial rearrangement ([8, 10, 16]). To map this conformational freedom across domains and ensure our analysis was statistically well-grounded, we clustered trajectory frames within regions of interest (ROIs) around each free-energy minimum (F1, F6 and T1) and sampled up to 50 representative structures per cluster to capture state heterogeneity (Supplementary Figs. S9–S10, Supplementary Table S9; see Methods).

Domain-wise, protomer-resolved RMSD distributions across these ROI-sampled structures show where conformational variability is concentrated (Fig. 3), while domain–domain centre-of-mass (COM) distances reveal the direction of the associated rearrangements (Fig. 4). Across F1 and F6, the kinase N-lobe, C-lobe and FATC remain comparatively conserved, as does FAT-TRD2, whereas the largest variations occur within the surrounding regulatory architecture. The PRD is particularly heterogeneous and strongly asymmetric between protomers: PRD-B remains within a relatively narrow RMSD range across the sampled states, whereas PRD-A shows substantially greater variability, reaching its largest values in F6 and T1. Thus, the major structural differences between the metastable ensembles are concentrated outside the conserved kinase core.

**Figure 3:**
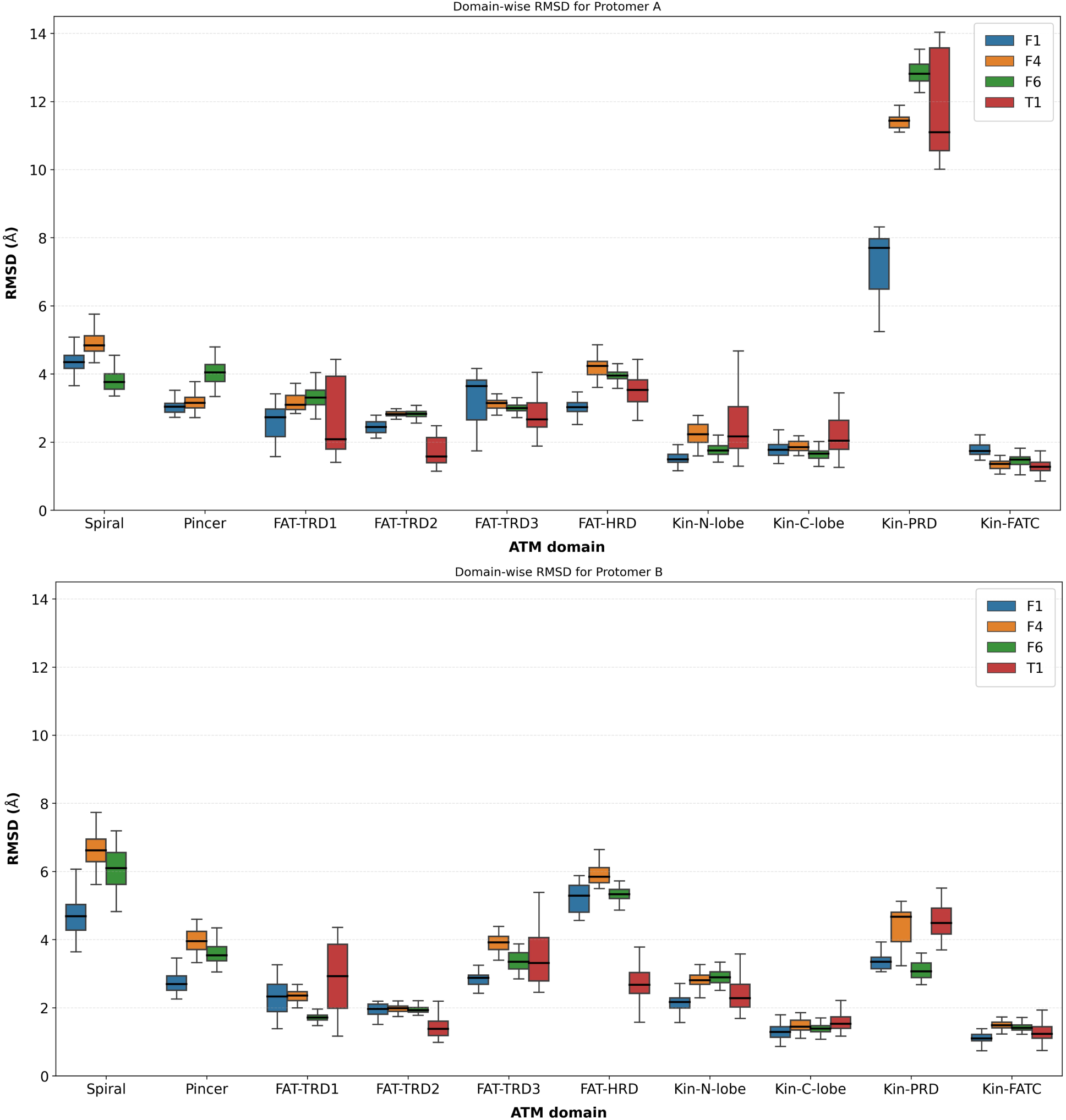
Domain-wise, protomer-resolved RMSD distributions across metastable substates. Backbone RMSD (relative to the post-equilibration reference structure; see Methods) for each ATM domain, computed from up to 50 representative structures per DBSCAN centroid within each substate ROI (see Methods) for the inactive (F1), alternative (F4), and oxidation-compatible (F6) full-length metastable states and the Spiral–Pincer-truncated global minimum (T1), shown separately for protomer A (top) and protomer B (bottom). Boxes show the interquartile range and median; whiskers extend to the most extreme non-outlier values. T1 lacks Spiral and Pincer domains by construction.

**Figure 4:**
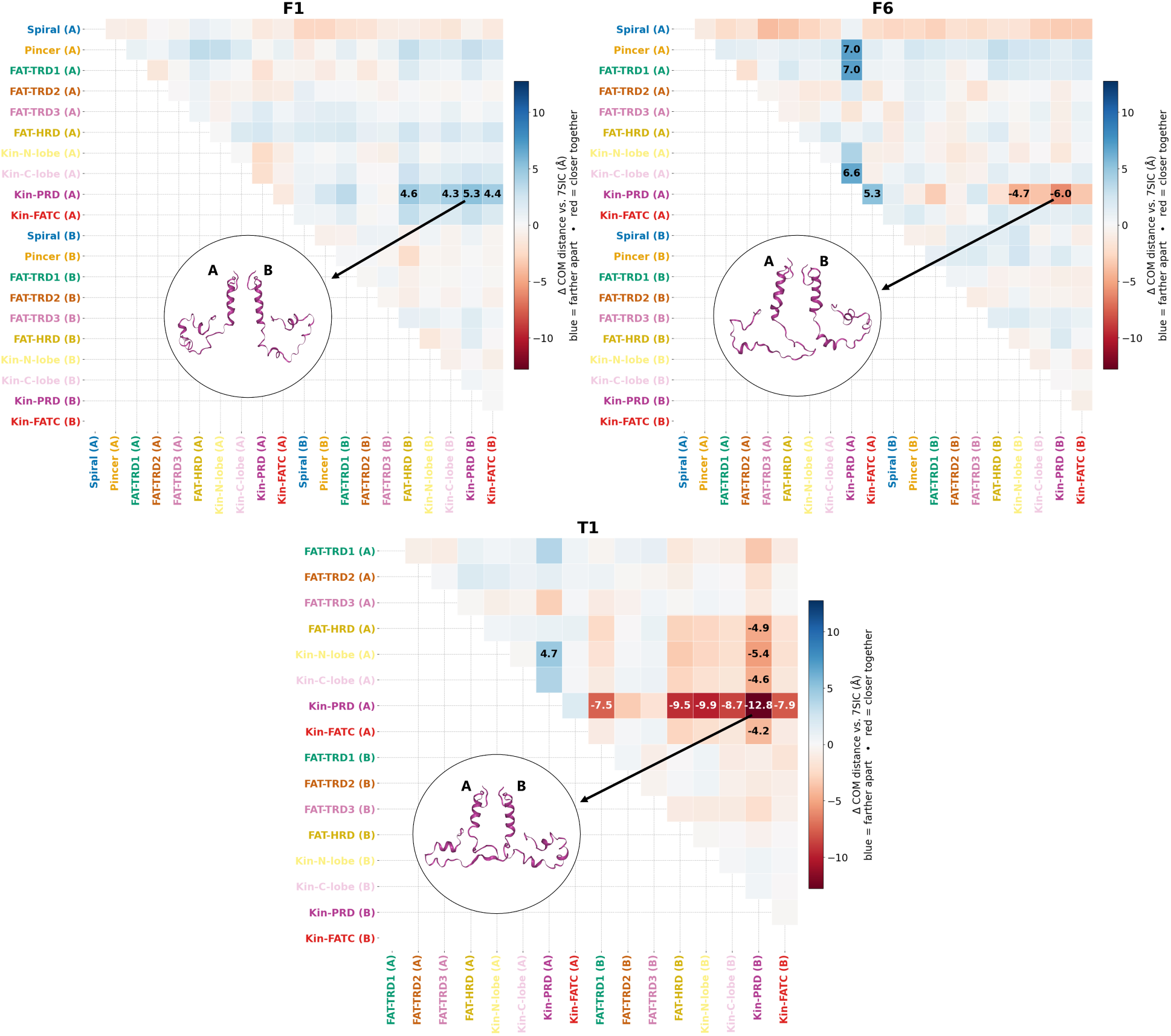
Domain–domain centre-of-mass (COM) distance differences relative to the experimental inactive structure (7SIC), frame-averaged over up to 50 representative structures per DBSCAN centroid within each ROI (see Methods). Each panel shows, for one metastable state (F1, F6 full-length; T1 truncated), the difference in inter-domain COM distance between that state and the experimental structure for every resolved domain pair.

The COM analysis further distinguishes the nature of these PRD rearrangements (Fig. 4). In F1, the elevated PRD-A RMSD is associated primarily with modest outward displacement relative to several domains of protomer B, consistent with the residual mobility of the inactive ensemble. In contrast, F6 shows a qualitatively different rearrangement: despite having the largest PRD-A RMSD, its COM distances indicate an inward movement, with PRD-A approaching PRD-B by approximately 6.0 Å and moving closer to the B-protomer kinase lobes by approximately 3.4–4.7 Å relative to experimental inactive structure (PDB: 7SIC). The large RMSD therefore reflects a directed repositioning of PRD-A rather than simply increased local disorder.

This inward PRD-A movement is amplified after Spiral–Pincer removal. In T1, PRD-A is closer to nearly all of the resolved domains than in 7SIC, with the largest change occurring toward PRD-B (−12.8 Å), more than twice the corresponding F6 displacement. Further, the rearrangement extends beyond the PRD itself: the broader RMSD distributions around the deleted scaffold include FAT-TRD1-A/B, the kinase lobes of protomer A and FAT-TRD3-B. Therefore, Spiral–Pincer removal does not simply increase flexibility uniformly across the dimer; instead, it redistributes conformational freedom into specific domain relationships, most prominently the PRD and its surrounding regulatory architecture.

Furthermore, an additional feature of F6 links this PRD repositioning to the distal scaffold. In the same protomer in which PRD-A moves inward, Spiral-A undergoes pronounced contraction relative to both neighbouring domains within protomer A and domains of protomer B. The coincidence of Spiral-A contraction with inward PRD-A movement suggests that the oxidation-compatible ensemble is associated with coordinated reorganization of the distal scaffold and PRD rather than an isolated fluctuation of the Cys2991-containing region.

### Experimental structures recapitulate domain repositioning and reveal protomer-specific PRD remodelling

To assess whether the simulated metastable states are consistent with experimentally observed ATM architectures, we compared the full-length inactive-like (F1) and oxidation-compatible (F6) ensembles, together with the Spiral–Pincer-truncated state (T1), with the experimentally determined inactive (PDB: 7SIC) [13] and oxidized (PDB: 8OXM) [8] structures. All comparisons were restricted to C*α* atoms of residues resolved in 7SIC, providing a common residue set across the experimental and simulated structures.

Structures were superposed using two complementary alignment schemes: alignment to the conserved FAT-TRD2 dimer interface (“TRD2-anchored RMSD”), which captures domain repositioning relative to the dimer core, and independent alignment of each domain (“independent RMSD”), which reports internal structural remodelling. Full domain-wise values are provided in Supplementary Table S8.

Comparison of F1 with the experimental inactive structure 7SIC shows that the simulated inactive-like state preserves the overall FAT–kinase architecture while exhibiting larger differences in the distal scaffold (Fig. 5A). The FAT and kinase core remain relatively close to 7SIC, with TRD2-anchored RMSDs of ≲ 5 Å and independent RMSDs of ≲ 2.7 Å, whereas Spiral and Pincer differ by approximately 12–16 and 6–10 Å, respectively. These differences are consistent with the intrinsic mobility of the N-terminal scaffold and do not compromise the inactive-like character of F1.

**Figure 5:**
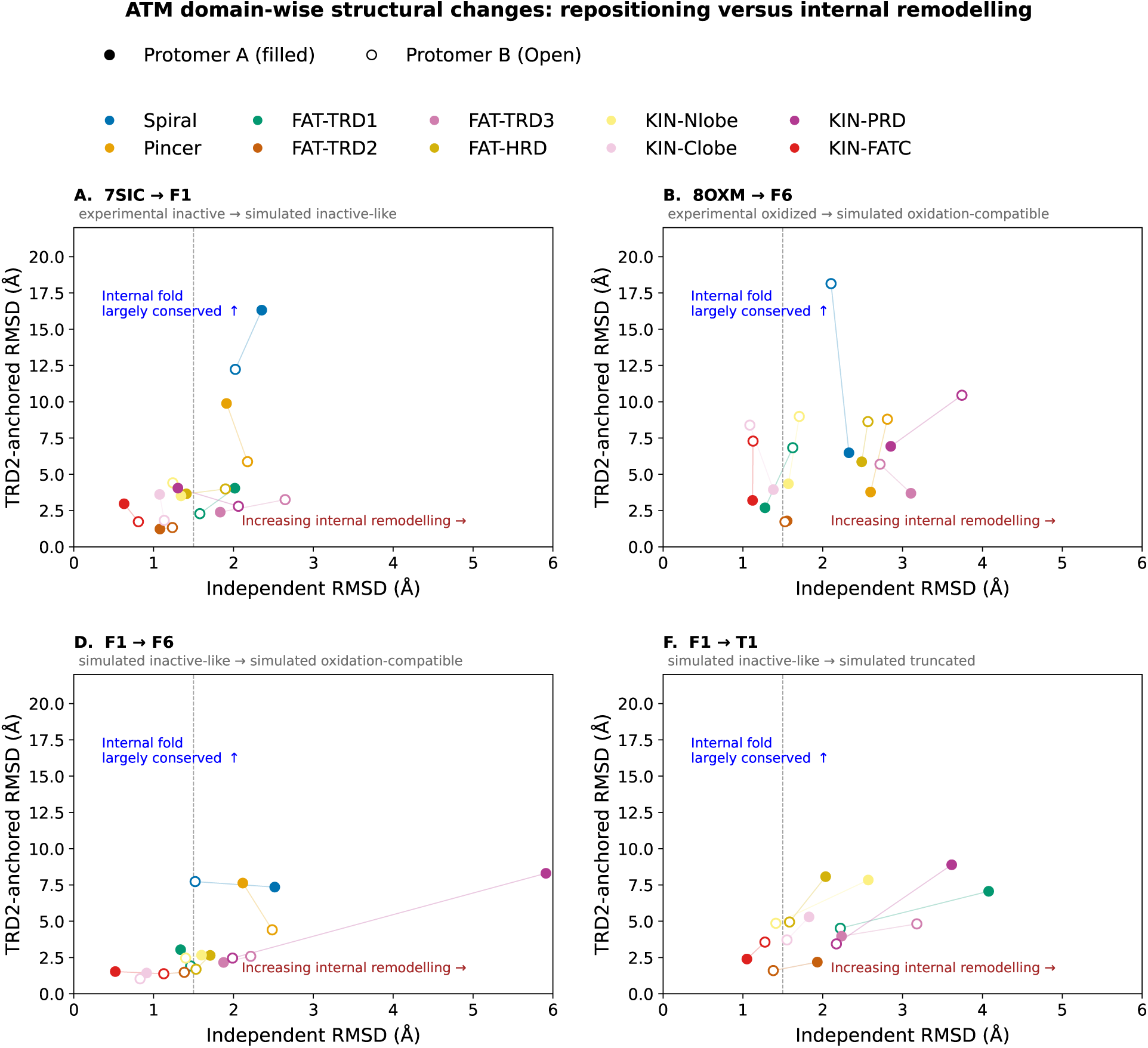
Domain repositioning and internal remodelling across experimental and simulated ATM states. Each panel compares domain-wise structural differences using independent RMSD (internal remodelling) and TRD2-anchored RMSD (domain repositioning relative to the conserved FAT–TRD2 dimer interface): (A) simulated inactive-like F1 versus experimental inactive ATM (7SIC), (B) simulated oxidation-compatible F6 versus experimental oxidized ATM (8OXM), (C) simulated inactive-like F1 versus oxidation-compatible F6, and (D) simulated inactive-like F1 versus Spiral–Pincer-truncated T1. Filled and open symbols denote protomers A and B, respectively. All RMSDs were calculated from C*α* atoms of residues resolved in 7SIC, providing a common residue set across the experimental and simulated structures. The plots distinguish domain repositioning from internal structural remodelling and highlight the pronounced, protomer-specific PRD remodelling in F6 and the broader redistribution of structural freedom following Spiral–Pincer removal.

The comparison of F6 with the experimentally oxidized structure 8OXM reveals a related but non-identical PRD arrangement (Fig. 5B). The independent PRD RMSDs are 2.9 Å and 3.7 Å for protomers A and B, respectively, indicating that F6 does not simply reproduce the experimentally determined oxidized structure. Because 8OXM was modelled by fitting the 7SIC structure into the activated maps [8], the comparison is interpreted primarily at the level of domain arrangement rather than as evidence for a unique internal structure. F6 therefore represents an alternative oxidation-compatible member of the ATM conformational ensemble. The PRD remodelling observed in F6 is also consistent with the activation-associated structural changes reported for constitutively active PRD-mutant ATM [9].

Direct comparison of the simulated inactive-like and oxidation-compatible states (F1*→*F6) shows that the transition involves selective remodelling of the regulatory architecture rather than wholesale reorganization of the protein (Fig. 5C). The conserved dimer framework and kinase core remain comparatively stable, whereas the largest changes are concentrated in the PRD and surrounding FAT–kinase regulatory elements. Consistent with the broader domain analysis, F6 is characterized by pronounced and protomer-specific PRD remodelling while retaining the overall dimeric framework.

In contrast, removal of the Spiral–Pincer scaffold produces a broader redistribution of structural freedom (Fig. 5D). Relative to F1, T1 shows increased internal remodelling across FAT-TRD1, FAT-TRD3 and the PRD, whereas FAT-TRD2 remains comparatively conserved. Thus, Spiral–Pincer removal affects not only the local geometry of the remaining PRD but also the organization of the surrounding FAT–kinase architecture. Together, these comparisons distinguish the selective, protomer-specific remodelling associated with the oxidation-compatible state from the broader structural redistribution caused by removal of the distal scaffold.

### Interaction-network remodeling accompanies protomer-specific PRD reorganization

We therefore next quantified interaction-network remodeling across F1, F6 and T1 using three complementary contact definitions: C*α*–C*α* proximity (*<* 8 Å) as a measure of geometric packing, hydrophobic side-chain contacts (*<* 5 Å), and acidic/basic residue contacts (*<* 4 Å) as a proxy for electrostatic interactions. Evaluating these metrics separately allows changes in domain packing to be distinguished from changes in specific chemical interactions. The analysis was performed both at the network level and for individual domain–domain interfaces (Fig. 6; Supplementary Fig. S11, S12, S13).

**Figure 6:**
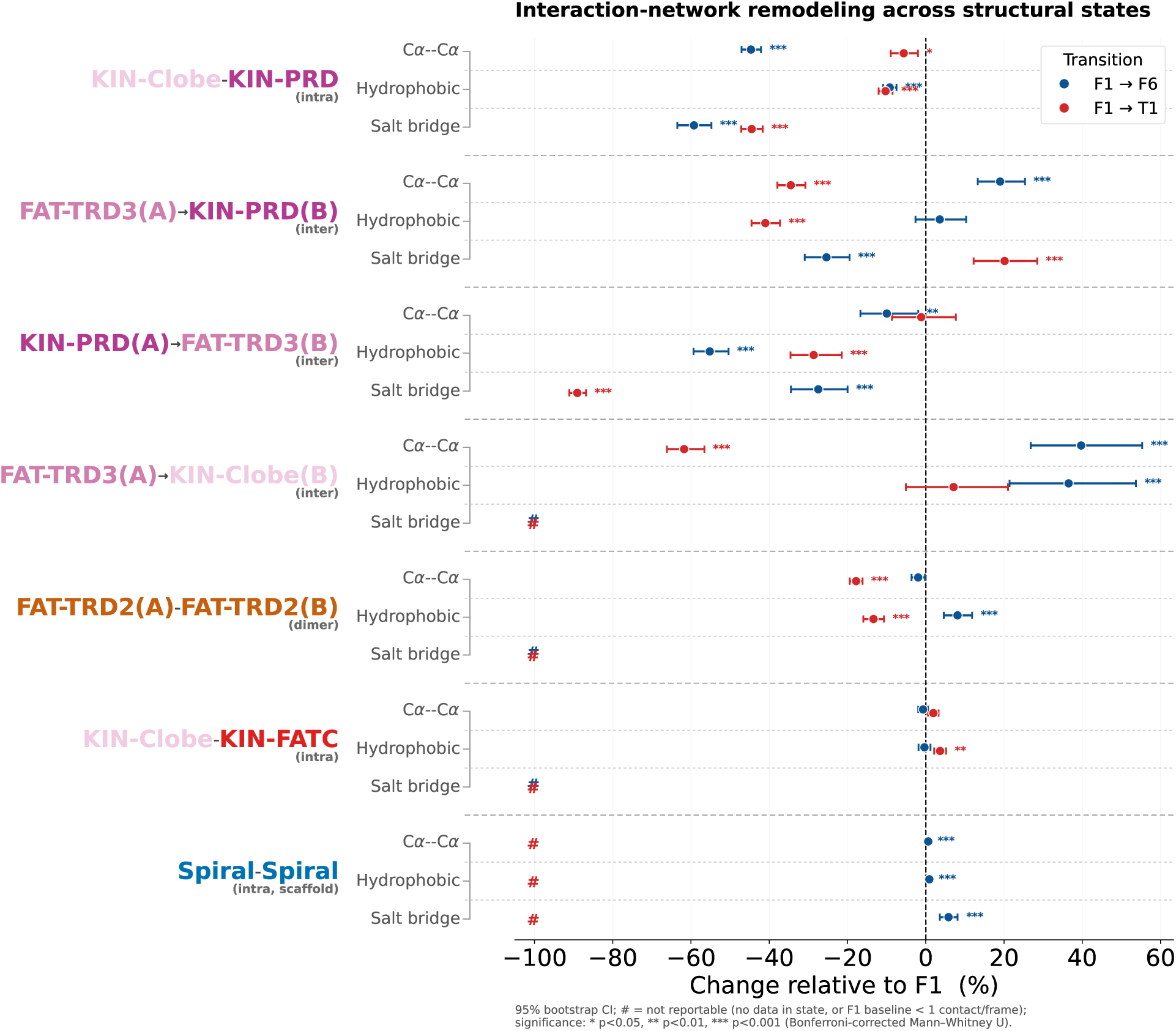
Quantification of interaction-network rewiring across structural states, for three contact metrics. For each of seven domain–domain interfaces, the percentage change in mean contact count per frame relative to F1 for the F1*→*F6 (blue) and F1*→*T1 (red) transitions, reported separately for C*α*–C*α* proximity (*<*8 Å), hydrophobic side-chain contacts (*<*5 Å) and acidic/basic residue contacts (*<*4 Å; contact definitions in Methods). For the salt-bridge metric, intra-protomer counts sum both ionic directions of each domain pair. Horizontal bars are 95% bootstrap confidence intervals (20,000 resamples); significance is assessed by two-sided Mann–Whitney U tests with Bonferroni correction (\**p <* 0.05, \*\**p <* 0.01, \*\*\**p <* 0.001). # marks non-reportable comparisons (no data in the state, or F1 baseline below one contact per frame). Interface labels are coloured by domain.

At the network level, F1-to-F6 remodeling is selective rather than a generalized loss of contacts. Inter-protomer connectivity decreases across the C*α*, hydrophobic and electrostatic networks, with the largest relative change occurring in the electrostatic component, whereas intra-protomer connectivity is comparatively preserved. T1 exhibits a broader reduction in inter-protomer C*α* and hydrophobic connectivity, consistent with the more extensive structural freedom observed after Spiral–Pincer removal. The distinction is therefore not simply between a connected and a disconnected dimer: F6 retains the overall interaction architecture while selectively redistributing inter-protomer coupling, whereas T1 produces a broader reorganization of the remaining network.

At the domain-pair level, the F1-to-F6 transition is concentrated around the FAT-TRD3–kinase–PRD region. The intra-protomer KIN-C-lobe–PRD C*α* contact decreases by approximately 45%, accompanied by increases in specific inter-protomer contacts involving FAT-TRD3. FAT-TRD3(A)–PRD(B) C*α* connectivity increases by approximately 19%, while FAT-TRD3(A)–KIN-C-lobe(B) connectivity increases by approximately 40%. The latter is accompanied by an approximately 37% increase in hydrophobic contacts. These changes are not uniform across interaction types: the hydrophobic contribution to FAT-TRD3(A)–PRD(B) is approximately unchanged, whereas its electrostatic component decreases by approximately 25%. Thus, the increased geometric proximity of selected domain pairs does not correspond to uniform strengthening of all contacts, but rather to a redistribution of the interaction modes supporting the reorganized PRD.

The dimeric framework remains comparatively conserved during F1-to-F6 remodeling. FAT-TRD2–FAT-TRD2 connectivity changes relatively little, and the Spiral–Spiral interface is also largely retained. The structural and network data therefore point to a localized reorganization within the FAT–kinase–PRD region while the principal dimeric scaffold remains intact. This provides a network-level explanation for how substantial PRD-A repositioning can occur without wholesale disruption of the ATM dimer.

The truncated T1 state provides a distinct perturbation. Removal of Spiral–Pincer produces broader changes in inter-protomer C*α* and hydrophobic connectivity and redistributes contacts toward the remaining FAT–kinase architecture. The FAT-TRD2–FAT-TRD2 interface is reduced relative to F1 but remains a major residual contact, while the inter-protomer FAT-TRD3–PRD and FAT-TRD3–KIN-C-lobe C*α* contacts are reduced. Thus, the broader structural divergence observed in T1 is accompanied by redistribution of the residual interaction network rather than complete collapse of the dimeric core.

The particularly strong state dependence of the electrostatic component raises a more specific question: which salt-bridge interactions account for the network-level changes, and are they distributed across the regulatory architecture or concentrated at selected domain interfaces?

### State-dependent reorganization of the salt-bridge network

Domain-resolved analysis of the same networks (Fig. 7) shows that the dominant state dependence lies in the electrostatic network. The cross-dimer FAT-TRD3–PRD salt-bridge interface identified in F1 progressively weakens through F6 and further declines in T1. Importantly, its protomer directionality is reorganized: the dominant contribution shifts from PRD-A–to–FAT-TRD3-B coupling in F1 to FAT-TRD3-A–to–PRD-B coupling in T1. Spiral–Pincer removal therefore does not produce a uniform loss of packing; rather, it selectively reorganizes electrostatic connectivity and inter-protomer coupling while preserving much of the intraprotomer interaction framework.

**Figure 7:**
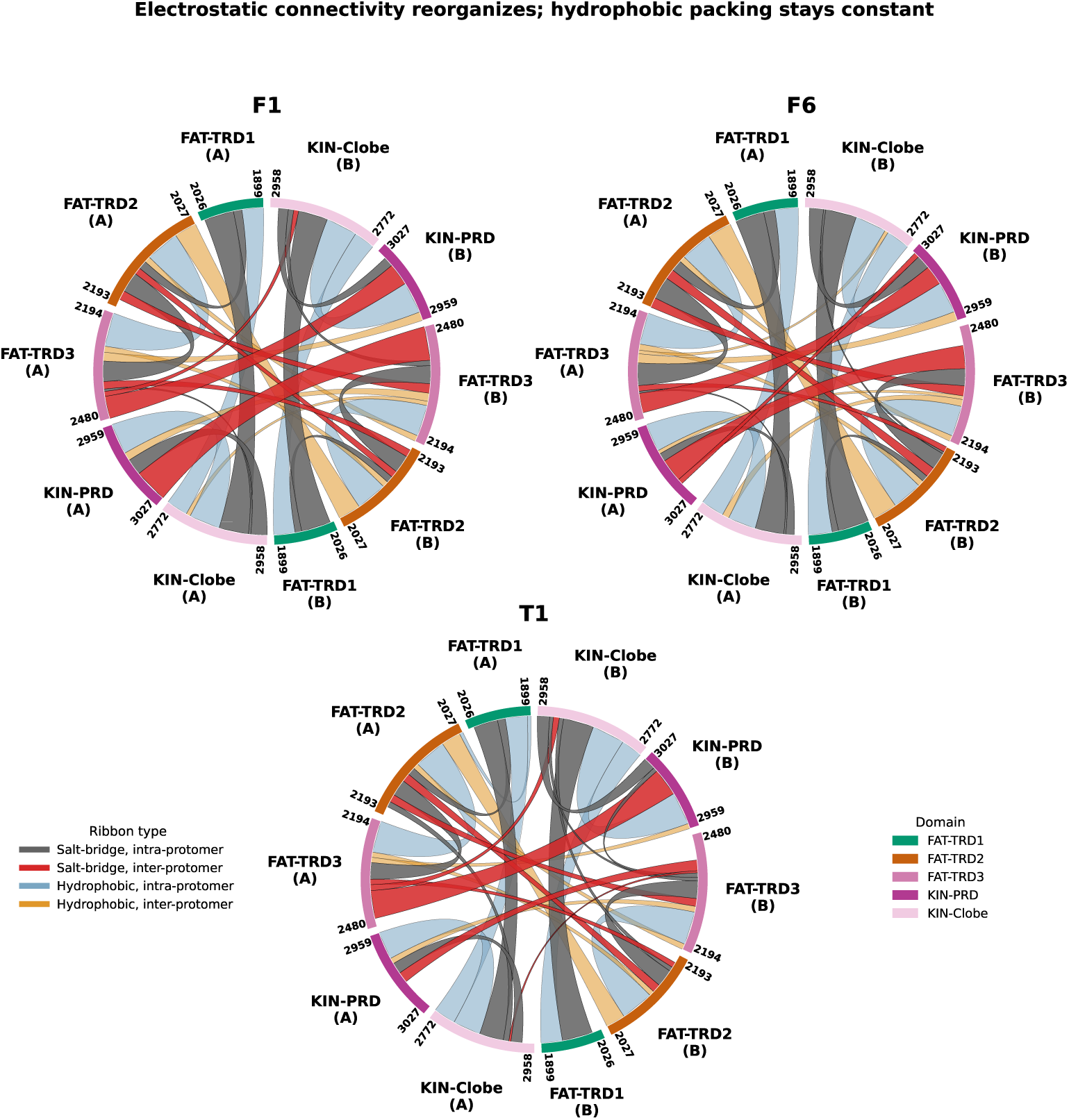
Domain- and residue-level reorganization of the salt-bridge and hydrophobic networks. Domain-level connectivity across F1, F6 and T1; grey/red ribbons show salt-bridge intra-/inter-protomer contacts, light blue/orange ribbons show hydrophobic intra-/inter-protomer contacts. Ribbon width is proportional to total occupancy; salt-bridge and hydrophobic ribbons use independently calibrated width scales owing to their differing raw magnitudes (see Methods).

To resolve this reorganization at the residue level, representative structures from the F1, F6 and T1 basins were subjected to short (5 ns) unbiased MD simulations. For each residue within the FAT–kinase–PRD architecture, salt-bridge degree (number of distinct interaction partners) and occupancy (the summed frame-fraction over a residue’s ionic contacts (Eq. (1)) were evaluated (Fig. 8; Methods). Across both transitions, degree changes relatively little for most residues, whereas occupancy exhibits greater variation. A small set of residues dominates both transitions. To identify the specific residue pairs underlying these changes, we further examined the contact frequencies of individual interactions (Supplementary Fig. S11). In F1*→*F6, the most prominent changes involve A:R2191 (FAT-TRD3), B:D2997 (PRD) and A:E3021 (C-terminal PRD), which gain occupancy with only modest changes in degree. In F1*→*T1, A:R2191 and A:E3021 instead show decreases in both degree and occupancy, whereas B:D2997 gains both. The increased connectivity of B:D2997 is associated with the formation of new inter-protomer salt bridges with A:K2418 and A:R2419 in FAT-TRD3, which are absent in F6. In F6, by contrast, B:D2997 forms an inter-protomer PRD–PRD contact with A:R2993, observed in 53.3% of sampled frames, which is absent in T1. These alternative interaction partners are associated with distinct PRD arrangements, and the resulting repositioning of the Cys2991-bearing loop could influence the geometry required for inter-protomer Cys2991–Cys2991 disulfide formation under oxidizing conditions.

**Figure 8:**
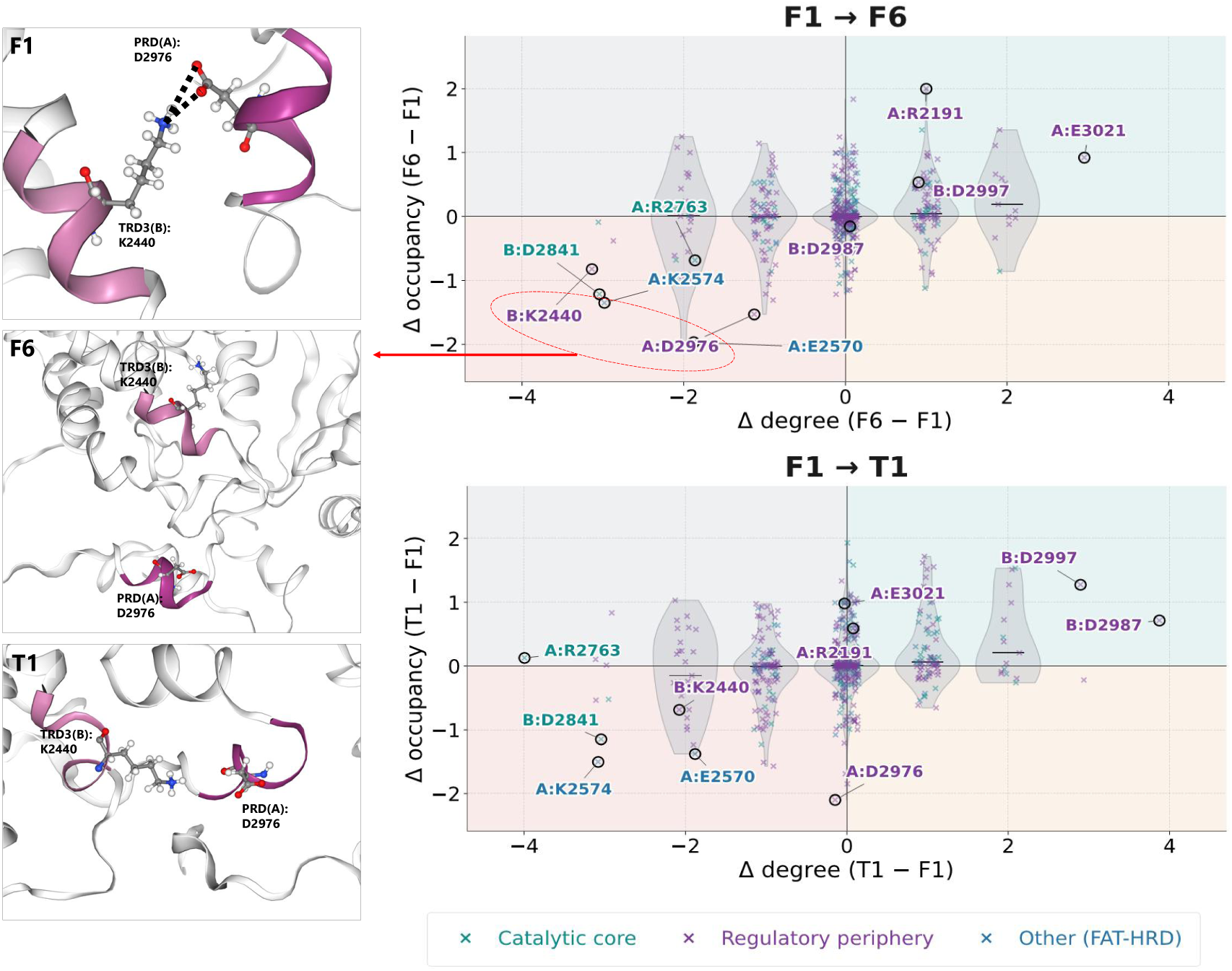
Residue-level salt-bridge reorganization across metastable states. Change in degree versus total occupancy for F1*→*F6 (top) and F1*→*T1 (bottom) for residues within the FAT, kinase, and PRD domains. Degree is the number of distinct partners; occupancy is summed contact persistence relative to F1. Shaded quadrants indicate change regimes, and violin plots summarize occupancy distributions per integer degree change. Points are colored by domain category with key residues labeled. (Insets) Cross-protomer B:K2440–A:D2976 salt bridge on k*α*9b, formed in F1 and lost in F6 and T1.

At the individual-contact level, A:E3021 shows a distinct pattern: it forms an intraprotomer salt bridge with A:R3012 in both F6 and T1, which is absent in F1, despite the reduction in its overall degree and occupancy in F1*→*T1. This reflects the loss of other interaction partners alongside the formation of a specific new contact, rather than a general increase in connectivity. Notably, this interaction occurs near K3016, an ATM acetylation site whose modification by Tip60 is required for DNA-damage-induced upregulation of ATM kinase activity [19].The state-dependent interaction pattern therefore identifies a region associated with an independently characterized regulatory modification.

A further contact links this rewiring to the autoinhibitory PRD helix k*α*9b and aligns with an experimentally validated site. Specifically, B:K2440 (FAT-TRD3) forms a high-occupancy salt bridge with A:D2976 on k*α*9b in F1 that is disrupted in both F6 and T1 (Fig. 8, insets), supporting the displacement of k*α*9b from its autoinhibitory posture during these transitions. Notably, D2976 is adjacent to Q2971, a residue whose mutation drives ATM hyperactivation [8]. By identifying a state-dependent salt bridge at a locus known to govern autoinhibition, our analysis reinforces the functional relevance of the recovered network.

### A mechanistic model for Spiral–Pincer-mediated restraint of ATM activation

Taken together, the free-energy, domain level, contact and residue level analyses converge on an integrated model of Spiral–Pincer function that departs from the conventional view of this *α*-solenoid scaffold as a passive MRN-docking platform or a conduit that actively transmits an activating signal to the kinase domain. Instead, the data support a restraint-release model. The intact Spiral–Pincer scaffold suppresses access to an oxidation-compatible conformational family that is already encoded within the intrinsic dynamics of the full-length dimer, and activation proceeds primarily by releasing this restraint – not by propagating a signal across the *∼*100 Å separating the N-terminal scaffold from the kinase active site.

Mechanistically, the available evidence is consistent with the following sequence.

i. The oxidation-compatible geometry (F6) exists as a thermally accessible, but energetically penalized, metastable state of the intact dimer, populated at 12.8 kcal mol*^−^*^1^ above the global minimum; this penalty is imposed by the Spiral–Pincer scaffold, since it collapses entirely upon its removal (Fig. 2b).
ii. Progression toward this state is not a concerted, symmetric rearrangement of the dimer. Instead it is led by protomer-specific remodelling of the PRD in one protomer – independently identified both from the electrostatic contact asymmetry of the PRD and from the domain-structural comparison against experimental structures, in which the PRD of protomer A alone shows a large *independent* RMSD in F6 (Supplementary Table S8).
iii. This asymmetric PRD remodelling occurs while the surrounding FATKIN architecture, and Cys2991 itself, remain largely disconnected from the Spiral–Pincer network at the residue level, indicating that the restraint acts on the PRD’s global positioning and conformational freedom rather than on Cys2991 directly.
iv. A newly formed inter-protomer KIN-PRD–KIN-PRD salt bridge (B:D2997–A:R2993, observed in F6, Fig. 8) and an intensified Spiral–Spiral inter-protomer contact accompany this transition, coupling the reorganization to both the upper *α*-solenoid interface and the lower FATKIN module simultaneously.
v. Removing the Spiral–Pincer scaffold does not simply remove this restraint uniformly. It collapses the associated free-energy penalty, intensifies the FAT-TRD3–KIN-PRD electrostatic interface beyond even the level seen in F6, and permits genuine internal structural remodelling of FAT-TRD1, FAT-TRD3, and the PRD – remodelling that is not observed anywhere in the intact, biologically relevant full-length landscape (Supplementary Table S8). This indicates that the scaffold constrains both the energetic accessibility and the structural fidelity of this regulatory region.

## Discussion

This study set out to resolve a question left open by every existing structural study of ATM: whether the long-range Spiral–Pincer scaffold merely correlates with kinase activation or causally regulates access to it, and how the disordered, experimentally unresolved Cys2991-containing PRD loop participates. Three independent lines of evidence converge on the same answer. The free-energy landscape shows the oxidation-compatible conformation is not generated *de novo* by Cys2991 oxidation, but selected from a pre-existing, energetically penalized metastable state of the native ensemble – a penalty imposed specifically by the Spiral–Pincer scaffold, not by the kinase regulatory domain itself. The salt-bridge network and the comparison against experimental structures independently identify the same asymmetry: activation-associated PRD remodelling is led by one protomer, not shared equally across the dimer, and the residue-level network indicates this restraint is transmitted through domain-scale coupling – not a direct residue-by-residue relay to Cys2991 – with kin C-lobe as a specific, testable relay point. Together, these results support a model in which ATM’s Spiral–Pincer scaffold acts as an allosteric restraint on the kinase regulatory module, whose release, not the propagation of a signal, licenses transition to a substrate-compatible state.

### Conformational selection as a general principle

The finding that the oxidation-compatible geometry pre-exists within the native ensemble situates ATM activation within the broader conformational-selection model of allosteric regulation. This agrees with the view that wild-type ATM exists in equilibrium between basal and active conformations, with the basal form strongly favoured [8, 9]. In this model, ligand or covalent triggers act by shifting the population of pre-existing conformational substates, rather than by inducing conformations absent from the unliganded ensemble [20–23]. This distinction matters mechanistically. A covalent trigger (Cys2991 oxidation) that merely stabilizes an existing minimum, rather than one that must first create a new conformational pathway, is a more parsimonious and kinetically favourable route to activation. This fits with the fact that ATM must be capable of activation on a timescale compatible with an acute genotoxic or oxidative insult.

### Spiral–Pincer as a restraint, not a conduit

The comparative analyses of systems with and without the Spiral–Pincer scaffold distinguish two pictures that are hard to separate from correlative structural data alone: a scaffold that actively transmits an activating signal, versus one that restrains access to an already-available activated state. A signal conduit whose removal abolishes or leaves unchanged the accessibility of the oxidation-compatible geometry is inconsistent with our results; instead, removal collapses its energetic penalty and permits additional remodelling not sampled in the intact protein – the signature of restraint release rather than loss of signal transmission. This reframes the Spiral–Pincer *α*-solenoid – conventionally described primarily as an MRN-docking platform [10, 13, 15] – as an element whose principal function may be to limit, by default, the conformational freedom of the distal kinase regulatory module. This interpretation is further supported at the residue level: the FAT-TRD1–kin C-lobe interface identified here as the strongest, truncation-responsive intra-domain contact corresponds structurally to one arm of the “C-clamp” previously described as stabilizing the inactive PIKK kinase-domain fold [13, 24], and its context-dependent strengthening upon loss of the restraining scaffold is consistent with a role in actively maintaining, rather than passively reflecting, the autoinhibited state. This interface lies close to the hinge between TRD1 and the FAT clamp, which rotates in activated-state structures [8, 9], so a restraint acting here would sit at a mechanically relevant point.

This model also provides an explanation for two observations that a simple signal-transmission picture does not. First, it explains why the oxidation-compatible state is rare, but not absent, in the resting full-length protein. A restraint that raises the free energy of a pre-existing conformational family, rather than an inactive state that lacks that family altogether, permits occasional, transient sampling of the oxidation-compatible geometry even under unstressed conditions. This provides a plausible structural basis for the low basal ATM activity observed in the absence of exogenous stimulation [8, 13], in marked contrast to the several-hundred-fold higher basal activity reported for *kα*9*b*-deleted or PRD-mutant ATM [8, 9]. Second, it explains why truncation does not simply abolish the transition. Instead, it makes the oxidation-compatible-like geometry the new global minimum while simultaneously permitting broader internal remodelling beyond that sampled in the full-length protein. Thus, removal of a restraint, rather than removal of an activating conduit, naturally produces both effects.

### A possible unifying logic for MRN-driven and oxidative activation

ATM’s two structurally distinct activation routes – MRN engagement following DNA double-strand breaks, and Cys2991 oxidation under oxidative stress [8, 13] – have generally been treated as mechanistically separate pathways converging only on downstream kinase activity. The restraint-release model suggests a more specific point of convergence. Both triggers may ultimately access a shared family of substrate-compatible ATM conformations through distinct molecular routes. This interpretation is consistent with recent structural evidence showing that PRD perturbation and oxidative activation are associated with globally similar active-state architectures, including coordinated PRD displacement, FAT-clamp rearrangement and kinase N-lobe/C-lobe realignment [9]. MRN-induced rearrangement of the N-terminal scaffold in one case, Cys2991 disulfide formation stabilizing a pre-existing PRD-proximal geometry in the other. This is a testable prediction: if correct, the metastable states populated transiently under MRN engagement and those populated under oxidative stress should show substantial structural overlap in the PRD/kinase-catalytic cleft region despite their distinct N-terminal triggers, a comparison not yet available from existing cryo-EM structures of the two activation modes and a natural target for future metadynamics or cryo-EM work.

These considerations also have implications for the response to ionizing radiation, which can engage ATM through DNA double-strand breaks and MRN while also promoting mitochondrial ROS and additional ATM activation [25, 26]. The existence of a pre-existing, oxidation-compatible conformational family therefore provides a structural framework for considering how DNA-damage and oxidative inputs might bias overlapping regions of the ATM conformational landscape, without requiring them to follow the same activation pathway.

### Generalization to the PIKK family

ATM shares its overall architecture, including the FAT– PRD–kinase cleft-occlusion logic, with mTOR, DNA-PKcs and the yeast orthologue Tel1, which are regulated by distal, non-catalytic-site inputs such as RHEB binding to mTOR and Ku–DNA engagement of DNA-PKcs [4, 14, 16, 27–30]. In mTORC1, RHEB binds approximately 60 Å from the kinase active site yet induces a *∼*3 Å realignment of active-site residues and increases *k*_cat_ by approximately 30-fold; cancer-associated hyperactivating mutations map to structural elements that maintain the inactive conformation and can therefore mimic relief of autoinhibition [31]. ATM shows a related principle: deletion or mutation of the autoinhibitory PRD helix (k*α*9b) produces MRN-independent hyperactivity and active-state structural rearrangements that are globally similar to those observed in oxidized ATM [8, 9]. In DNA-PKcs, cryo-EM structures captured a sequential transition from inactive to activated states in which Ku–DNA engagement is accompanied by stretching and twisting of HEAT-repeat solenoids, with these long-range rearrangements propagated to the FAT–kinase module and coupled to kinase activation [32]. Thus, across several PIKKs, distal regulatory elements can control kinase activation through large-scale rearrangement of the surrounding architecture rather than by directly remodeling the catalytic site. These observations, however, do not by themselves distinguish between selection of pre-existing active conformations and induced conformational change, nor do they establish whether the distal element functions primarily as a restraint or as a signal-transmission conduit. Our results address these questions specifically for ATM by showing that the Spiral–Pincer scaffold restrains access to a pre-existing oxidation-compatible conformational family. If this restraint-release principle proves to be a broader feature of PIKK regulation, ATM would extend it to a chemically distinct covalent trigger, in which Cys2991 oxidation stabilizes an otherwise rare member of the conformational ensemble. Testing this possibility in ATM’s MRN-driven pathway, or through analogous perturbation and free-energy analyses of other PIKKs such as mTOR or Tel1, will be required to determine its generality.

### Relevance to disease and therapeutics

ATM inhibitors are being explored clinically as radiosensitisers, while loss-of-function ATM mutations cause ataxia-telangiectasia [2, 33]. Most current inhibitors target the ATP-binding site directly. The restraint-release mechanism identified here suggests a complementary, non-catalytic strategy: stabilizing the Spiral–Pincer-constrained architecture could in principle suppress the conformational transition required for activation, with the FAT-TRD1–kin C-lobe and Spiral–Spiral interfaces as candidate sites. Conversely, disrupting or mimicking restraint release could enhance ATM activity where residual function is therapeutically desirable. Both possibilities remain speculative – we identify structural and energetic features of restraint release, not ligandability – and would require experimental follow-up.

In summary, we developed an atomistic model of the full-length human ATM dimer (*∼*3,056 residues), combining experimentally resolved structure with AlphaFold3-based completion of unresolved regions, including the Cys2991-containing PRD, and characterized its conformational land-scape using well-tempered metadynamics. Comparison with a Spiral–Pincer-truncated construct allowed us to isolate the contribution of the distal scaffold to ATM conformational regulation. Our simulations reveal a pre-existing pathway to an oxidation-compatible conformation in the native dimer, with the Spiral–Pincer scaffold acting as an energetic restraint on access to this state. Its removal shifts the oxidation-compatible-like geometry towards the lowest-energy state while releasing broader, protomer-asymmetric remodelling across the FAT–kinase–PRD regulatory architecture, without substantial deformation of the catalytic core. Together with the candidate FAT-TRD1–kin C-lobe relay and FAT-TRD3–PRD interaction changes, these findings provide a mechanistic and experimentally testable framework in which Spiral–Pincer regulates ATM activation by restraining, rather than transmitting, access to a pre-existing substrate-compatible conformational state.

## Methods

### Construction of full-length and truncated ATM structural models

The complete FASTA sequence of the human ATM homodimer was provided to AlphaFold3 (AF3) [34, 35] as two protein chains, using default model parameters and a fixed random seed, to generate structural models of the full-length dimer for use as the starting point for the MD and metadynamics simulations described below. The top five AF3 models were screened against the experimental inactive human ATM structure (PDB 7SIC, 2.53 Å resolution [13]) by global and domain-wise structural alignment and by comparison of AF3 confidence metrics (pTM, ipTM, per-domain pLDDT). Final model (Model 1) was selected as the representative structure for all subsequent simulations on the basis of its substantially closer global and domain-wise agreement with the experimental structure relative to the other four models, while showing no atomic clashes and a low fraction of predicted disordered residues. Notably, because both the PRD and the FAT-TRD3 domain of the opposing protomer have unresolved residues at their mutual interface in the experimental structure, AF3 reconstruction of both regions in Model 1 directly visualizes the proposed FAT-TRD3–PRD substrate-occlusion interaction [8, 10] for the first time, providing structural motivation for the free-energy and salt-bridge analyses of this interface reported in the Results. The Spiral–Pincer-truncated model was constructed by removing the corresponding residue range from Model 1 and capping and relaxing the resulting termini prior to solvation (see Molecular dynamics simulation protocol, below). Full details of the AF3 modelling procedure, model screening and selection criteria, confidence metrics, and domain-by-domain structural assessment of Model 1 are provided in Supplementary Note 1.

### Molecular dynamics simulation protocol

The full-length ATM homodimer comprised 3,056 residues per protomer (6,112 protein residues in total) and was solvated with 529,393 water molecules and 44 Na^+^ ions, yielding a system of 1,687,051 atoms. The Spiral–Pincer-truncated homodimer comprised 1,157 residues per protomer (2,314 protein residues in total) and was solvated with 114,702 water molecules and 10 Na^+^ ions, yielding a system of 381,606 atoms. Unbiased molecular dynamics (MD) simulations of both models were performed using GROMACS (version 2024.2) [36], the AMBER99SB-ILDN force field [37] and the TIP3P water model. Each protein model was solvated in a cubic box with a minimum distance of 1.0 nm between the protein and the box boundaries, and the system was neutralized with Na^+^ ions. Following steepest-descent energy minimization, the system was equilibrated for 5 ns in the NVT ensemble with positional restraints on the protein, followed by 5 ns of unrestrained NPT equilibration at 300 K and 1 bar. Further, production MD simulations were performed using a 2 fs timestep for 10 ns. Temperature and pressure were maintained using a velocity-rescaling thermostat (*τ* = 0.1 ps) and Parrinello–Rahman barostat (*τ* = 2 ps), respectively. Long-range electrostatic interactions were calculated using particle-mesh Ewald (PME), with 1.0-nm cutoffs for short-range electrostatic and van der Waals interactions. Bonds involving hydrogen atoms were constrained using LINCS, and periodic boundary conditions were applied in all three dimensions.

### Well-tempered metadynamics simulations

Given the computational cost of well-tempered metadynamics for systems of this scale (approximately 1.69 million atoms for full-length ATM and 0.38 million atoms for the Spiral–Pincer-truncated construct), computational effort was prioritized toward extended sampling (till convergence reached), with one trajectory performed per construct rather than multiple independent replicas. Convergence was assessed using within-trajectory diagnostics, including progressive decay of the deposited bias, stability of the reconstructed free-energy surface over the final 100 ns, and convergence of the relative free energies of all identified metastable states. WTMetaD simulations were performed with GROMACS 2024.2/PLUMED 2.9.2. Two collective variables (CVs) were simultaneously biased: the inter-protomer Cys2991 sulfur–sulfur distance (CV1) and the distance between the centre of mass of the PRD helix (k*α*9b) and the activation-loop residues within the same protomer (CV2). Gaussian potentials with an initial height of 4.18 kJ mol*^−^*^1^ and width of 0.35 nm were deposited every 2 ps using a bias factor of 15 at 300 K. The full-length and truncated systems were simulated for 500 ns and 196 ns ensuring convergence, respectively.

#### Rationale for collective variable selection

The efficiency and interpretability of metadynamics depend critically on selecting CVs that distinguish the relevant metastable states and transition regions; inadequate CVs can yield unreliable free-energy estimates and misleading transition mechanisms [21, 22]. We selected two physically motivated, low-dimensional distance CVs to monitor distinct structural features associated with the biological question: inter-protomer Cys2991 proximity and PRD displacement relative to the substrate-binding cleft.

First, CV1 is the inter-protomer sulfur–sulfur distance between the two Cys2991 residues. Formation of an intermolecular Cys2991 disulfide bond during oxidative ATM activation requires these residues to approach one another [7, 8]. The sulfur–sulfur distance therefore provides a direct, physically interpretable measure of this geometric prerequisite, without relying on an indirect proxy that would require subsequent validation against cysteine proximity.

Second, CV2 is the distance between the centre of mass of the PRD helix (*kα*9*b*) and the activation-loop residues lining the substrate-binding groove within the same protomer. It was chosen to monitor the relative positioning of the PRD and the substrate-binding cleft, providing a low-dimensional geometric proxy for PRD-mediated cleft occlusion. The biological relevance of this coordinate is supported by recent structural and biochemical evidence showing that deletion of *kα*9*b* produces constitutive ATM activation and permits an active-state conformation with substrate bound under ligand conditions in which wild-type ATM remains in the basal conformation [9]. Thus, the PRD–activation-loop separation provides a physically interpretable measure of the relative positioning of these regulatory elements, although it does not directly quantify substrate accessibility or capture all aspects of their relative orientation. Related precedent comes from kinase metadynamics studies: Sutto and Gervasio used CVs based on selected atomic distances and contact-map differences relative to active and inactive activation-loop conformations to characterize the conformational free-energy landscape of EGFR kinase [38]. Because ATM shares the PIKK/PI3K-kinase-like fold and associated substrate-cleft regulatory architecture [10, 14], monitoring PRD–activation-loop separation is a mechanistically motivated choice.

Beside this, using two low-dimensional, physically interpretable distances keeps the resulting FES directly interpretable in terms of the two structural features motivating this study: Cys2991 proximity and PRD positioning relative to the substrate-binding cleft. This is consistent with the principle that CVs should be as few as practicable while still distinguishing the metastable states of interest [22]. This dimensional reduction necessarily coarse-grains over other potentially relevant degrees of freedom, including Spiral–Pincer orientation, PRD rearrangements beyond the selected distance, and local side-chain rotamers governing the Cys2991 disulfide-forming geometry.

### Free-energy surface reconstruction and identification of conformational states

The complete WTMetaD trajectories were used to reconstruct two-dimensional free-energy surfaces (FESs) as functions of CV1 and CV2. Trajectory reweighting accounted for the time-dependent metadynamics bias, and the resulting probability distributions were converted to free energies. Each FES was normalized by setting its global minimum to zero. Local and global minima were identified using watershed segmentation [39], implemented through the scikit-image library in Python; additional features of interest were identified by visual inspection. Because watershed-derived minima need not coincide exactly with grid points in intermediate FESs, free-energy values were sampled within a local region of interest (ROI) extending 0.3 nm around each reference minimum in the two-dimensional CV space. This neighbourhood is comparable in scale to the Gaussian bias width of 0.35 nm and accommodates modest shifts in the apparent minimum position while retaining a localized estimate of the corresponding basin free energy. Representative structures were then obtained using density-based spatial clustering of applications with noise (DBSCAN), enabling characterization of the conformational ensembles associated with the identified FES features (full sub-cluster population counts and their proportional sampling weights are given in Supplementary Table S9).

#### Convergence assessment of well-tempered metadynamics simulations

Convergence was assessed using three complementary within-trajectory diagnostics: the evolution of deposited Gaussian hill heights, the stability of the reconstructed free-energy surface (FES), (Supplementary Fig. S7) and the stability of relative free energies of predefined metastable states (Supplementary Fig. S8).

Deposited hill heights decreased overall in both systems (Supplementary Fig. S7a,d), remaining low during approximately the final 130 ns for the truncated model. The full-length model exhibited a similar trend, although occasional late increases suggested continued exploration of less-biased regions of collective-variable (CV) space. Hill-height evolution was therefore treated as a qualitative indicator of progressive bias saturation, rather than as a standalone criterion for convergence.

To assess FES stability, the final two-dimensional landscapes were analysed by visual inspection and watershed segmentation to identify the major minima. These minima defined a reference set for the subsequent analysis. Intermediate FESs were reconstructed every 1000 deposited Gaussian hills (2 ns) and normalized to their respective global minima before comparison along CV1 and CV2 (Supplementary Fig. S8). The close overlap of the profiles reconstructed over the final 100 ns (Supplementary Fig. S7b,c,e,f) supports stabilization of the major features of the sampled freeenergy landscapes.

Finally, the relative free energies of the predefined metastable states were tracked across successive FES reconstructions (Supplementary Fig. S8). For each intermediate FES, the relative free energy of each state was calculated with respect to the global minimum of that FES. Stabilization of these relative free-energy differences, together with persistence of the corresponding basins across successive reconstructions, was used to ensure convergence of the identified metastable states.

Taken together, these diagnostics support within-trajectory stabilization of the reconstructed FES and the relative free energies of the identified metastable states.

### Structural characterization of metastable ensembles

Structural analyses were performed on representative structures extracted from the identified metastable-state and transition-state ROIs. Domain-wise backbone RMSDs were calculated independently for each major ATM domain, using the final frame of production run as the reference for the corresponding model. This analysis distinguished local conformational changes within individual domains from larger-scale rearrangements of the protein. Changes in quaternary organization were quantified using COM distances between selected domains across the representative structures associated with different FES minima and ROIs, providing measures of relative domain positioning and compactness.

To assess whether Spiral–Pincer removal alters the local structure of the catalytic machinery, backbone and side-chain dihedral angles were calculated for selected kinase-domain residues and compared across representative structures from different FES regions. Secondary-structure assignments were obtained using DSSP, with particular attention to structural elements within the kinase domain and PRD. Distances between selected residues or structural elements previously implicated in ATM regulation were also calculated to characterize state-dependent changes in their relative positioning. The residue boundaries used to define the Spiral, Pincer, FAT-TRD1/2/3, FAT-HRD and kinase N-lobe, C-lobe, PRD and FATC regions are provided in Supplementary Table S10 and are consistent with the domain definitions used in Fig. 2 and the interaction-network analyses.

Following identification of the metastable basins, representative cluster-centroid structures from the F1 and F6 regions of interest (ROIs) in full-length ATM and the T1 ROI in truncated ATM were subjected to 5-ns unbiased molecular dynamics simulations. Simulations were performed using GROMACS with the AMBER99SB-ILDN force field, a 2-fs integration time step, and periodic boundary conditions. The systems were maintained at 300 K using a velocity-rescaling thermostat and at 1 bar using a Parrinello–Rahman barostat. Long-range electrostatic interactions were treated using the particle-mesh Ewald (PME) method, with 1.0-nm cutoffs for short-range electrostatic and van der Waals interactions. Hydrogen-involving bonds were constrained using the LINCS algorithm, and the initial velocities were retained without regeneration. Trajectory coordinates were recorded every 1 ps for subsequent structural/residue interaction analysis.

### Interaction and contact network analysis

Inter- and intra-protomer contacts were analysed across representative frames from the identified metastable ensembles to characterize state-dependent changes in ATM packing and interaction networks. Three complementary contact definitions were used to distinguish coarse-grained structural organization, hydrophobic packing and electrostatic interactions.

Backbone-level packing was characterized using C*α*–C*α* distances. Two residues were considered to be in contact when the distance between their C*α* atoms was *≤* 8 Å. This criterion provides a coarse-grained measure of residue proximity and domain packing without implying a specific chemical interaction.

Hydrophobic contacts were defined between side-chain carbon atoms of hydrophobic residues (A, V, I, L, M, F, W, Y and C), excluding C*α* atoms. Two hydrophobic residues were considered to be in contact when the distance between any pair of their side-chain carbon atoms was *≤* 5 Å. This definition was used to characterize state-dependent changes in non-polar packing.

Electrostatic contacts were identified using distances between selected positively and negatively charged side-chain atoms. K NZ and R NH1/NH2 atoms were defined as positively charged centres, whereas D OD1/OD2 and E OE1/OE2 atoms were defined as negatively charged centres. Oppositely charged atom pairs separated by *≤* 4 Å were classified as ionic contacts. Within each frame, multiple atom-level contacts involving the same residue pair were collapsed into a single residue-level interaction, preventing repeated counting of the same residue pair. These distance-based contacts provide a geometric representation of potential ionic interactions rather than a direct measure of interaction energy. The contact-metric comparisons at the network and domain-pair level (Fig. 6, Fig. 7) were computed from the DBSCAN-sampled representative structures of the metadynamics-derived metastable ensembles (F1, F6, T1), which provide broad conformational coverage of each basin for aggregated, relative comparisons.

### ROI-based ionic-contact network analysis

Unlike aggregated metrics, residue-level degrees and occupancies (Fig. 8, Supplementary Fig. S11) were computed from short (5 ns) unbiased MD trajectories to avoid bias-induced distortions of contact lifetimes inherent to metadynamics. Matching each analysis to its resolution—robust domain trends from the metastable ensemble and frame-wise occupancies from unbiased dynamics—yields consistent, mutually reinforcing insights, such as the shared directional reorganization of the FAT-TRD3–PRD interface. Residues were uniquely identified by protomer, residue number and amino-acid identity, allowing intra- and inter-protomer interactions to be distinguished. Each unique residue pair was represented by a single undirected edge, with its weight defined by the number of frames in which the contact persisted during the corresponding unbiased MD simulation.

Networks were constructed using NetworkX[40], with residues represented as nodes and ionic-contact pairs as edges. Two network measures were calculated for each residue. Degree (*k_i_*) was defined according to the NetworkX convention as the number of edges incident on node *i*, corresponding here to the number of distinct residues with which it formed ionic contacts, irrespective of edge weight.

Occupancy (*O_i_*) was defined as the sum of contact frequencies across all salt-bridge partners of residue *i*:

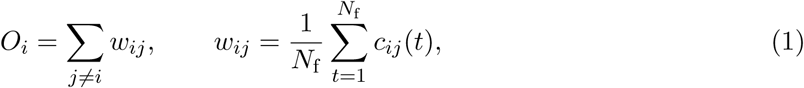

where *c_ij_*(*t*) = 1 if an ionic contact is present in frame *t* and 0 otherwise, and *N*_f_ is the total number of frames in the unbiased MD trajectory.

Thus, degree quantifies the diversity of a residue’s interaction partners, whereas occupancy represents the cumulative frame-wise occurrence of its ionic contacts across the sampled trajectory. Both measures were evaluated independently for F1, F6 and T1 to characterize state-dependent differences in residue connectivity and ionic-contact organization.

## Data availability

All configuration, structure, and run files related to the molecular dynamics (MD) and metadynamics simulations in this manuscript are publicly available on Zenodo (https://doi.org/10.5281/zenodo.23413684) under a Creative Commons Attribution 4.0 International (CC BY 4.0) licence. The deposited dataset includes AlphaFold-predicted structural models of full-length and truncated ATM, the experimental basal ATM crystal structure (PDB ID: 7SIC), and representative structures of the regions of interest (ROIs)—specifically F1 and F6 for full-length ATM, and T1 for truncated ATM. Additionally, the repository contains all input files required to reproduce the simulations (topology, equilibration, and production MD files), alongside metadynamics outputs including free energy surfaces, collective variables, hills files, and 1000-frame trajectory videos for both systems.

## Acknowledgements

The authors extend their thanks to Shri Probal Chaudhury (Associate Director, HS & E Group), Dr. S. Anand, HPD, and Mr R K B Yadav, RSSD, for their unwavering support and encouragement throughout the project. The authors also thank Mr. P P Shete, Computer Division, BARC for his steadfast support in the local installation and use of AlphaFold3.

## Author contributions

Conceptualization, A.C. and B.S.; methodology, A.C. and B.S.; software, A.C.; validation, A.C.; formal analysis, A.C.; investigation, B.S.; resources, B.S.; data curation, A.C.; writing—original draft preparation, A.C. and B.S.; writing—review and editing, B.S.; visualization, A.C.; supervision, B.S.; project administration, B.S. All authors have read and agreed to the published version of the manuscript.

