## Supporting information for "Long-range allosteric restraint by the Spiral–Pincer scaffold gates oxidative activation of the ATM kinase"

#### Contents

**Supplementary Note 1: AlphaFold3 model construction and validation** **2**

**Supplementary Tables** **13**

**Supplementary Figures** **21**

### Supplementary Note 1: AlphaFold3 model construction and validation

#### Rationale and input

The human ATM protein is a large (6112-residue) homodimeric serine/threonine kinase whose size and conformational complexity make complete high-resolution experimental structural data difficult to obtain. We used AlphaFold3 (AF3) [1] to generate structural models of the full-length human ATM dimer for use as the starting point for the molecular dynamics and metadynamics simulations described in the main text. The complete FASTA sequence of the human ATM homodimer was provided to AF3 as two protein chains, with no ion or ligand supplied as additional input. Default AF3 model parameters were used, with the random seed set to 1.

#### Completeness of the experimental reference structure

The best available experimental human ATM structure (PDB 7SIC, 2.53 Å resolution [2]) has several regions with missing (unresolved) residues, summarized by domain in Table S1. The PRD, Spiral, and FAT-HRD domains show the largest fractions of missing residues (39%, 15%, and 12%, respectively); the authors of the deposited structure attribute this to conformational flexibility in these regions, consistent with their absence from density in the cryo-EM map [2].

Table S1: **Residue completeness of the experimental ATM structure (PDB 7SIC) by domain.**

| Domain | Experimentally confirmed residues | Expected residue count | Missing residues (%) |
|---|---|---|---|
| Spiral | 994 | 1166 | 15% |
| Pincer | 696 | 734 | 5% |
| FAT-TRD1 | 116 | 125 | 7% |
| FAT-TRD2 | 157 | 165 | 5% |
| FAT-TRD3 | 273 | 286 | 5% |
| FAT-HRD | 120 | 137 | 12% |
| KD-Nlobe | 156 | 156 | 0% |
| KD-Clobe | 187 | 187 | 0% |
| KD-PRD | 40 | 66 | 39% |
| KD-FATC | 34 | 34 | 0% |

#### Sequence composition

Across the ATM dimer, 39% of residues are hydrophobic/non-polar, 25% are polar, 14% are polar and basic, 13% are polar and acidic, 5% are hydrophobic/non-polar and aromatic, and 3% are polar and aromatic. Hydrophobic/non-polar and polar residues dominate most domains (Table S2 and Table S3). The high hydrophobic content of the FAT and kinase domains is consistent with these regions forming compact structural cores, which likely explains why the kinase domain is comparatively well-resolved and accurately predicted by AF3. The polar/basic and polar/acidic residues together make up 27% of the sequence, consistent with the high charge content expected for catalysis, ATP binding, and substrate recognition. The PRD and FAT-TRD3 domains show comparatively low hydrophobic and high polar/charged content, consistent with their observed flexibility, missing density in the cryo-EM structure, and lower AF3 confidence (below), and consistent with the possibility that the FAT-TRD3–PRD interaction discussed in the main text, while important for substrate-site occlusion, involves transient or flexible contacts rather than a single rigid interface.

Table S2: **Domain-wise distribution of residue chemical type (residue counts, single protomer).**

| Residue type | Spiral | Pincer | TRD1 | TRD2 | TRD3 | HRD | Nlobe | Clobe | PRD | FATC |
|---|---|---|---|---|---|---|---|---|---|---|
| Hydrophobic/non-polar | 447 | 296 | 47 | 61 | 104 | 59 | 67 | 77 | 23 | 19 |
| Hydrophobic/non-polar, aromatic | 61 | 43 | 5 | 7 | 13 | 7 | 5 | 14 | 3 | 3 |
| Polar | 313 | 183 | 31 | 41 | 73 | 29 | 39 | 40 | 15 | 8 |
| Polar, acidic | 141 | 91 | 19 | 24 | 40 | 15 | 18 | 26 | 14 | 1 |
| Polar, aromatic | 32 | 26 | 6 | 11 | 9 | 2 | 3 | 6 | 1 | 0 |
| Polar, basic | 166 | 101 | 17 | 21 | 47 | 25 | 24 | 24 | 10 | 3 |

Table S3: **Domain-wise amino-acid composition (residue counts, single protomer).**

| Residue | Spiral | Pincer | TRD1 | TRD2 | TRD3 | HRD | Nlobe | Clobe | PRD | FATC |
|---|---|---|---|---|---|---|---|---|---|---|
| ALA | 57 | 32 | 10 | 10 | 20 | 8 | 8 | 7 | 3 | 2 |
| ARG | 56 | 24 | 6 | 9 | 17 | 13 | 11 | 11 | 4 | 1 |
| ASN | 57 | 32 | 2 | 4 | 14 | 8 | 7 | 7 | 4 | 2 |
| ASP | 54 | 46 | 9 | 5 | 10 | 6 | 10 | 9 | 6 | 1 |
| CYS | 39 | 23 | 2 | 4 | 8 | 2 | 4 | 6 | 1 | 0 |
| GLN | 59 | 31 | 6 | 8 | 18 | 3 | 12 | 9 | 5 | 3 |
| GLU | 87 | 45 | 10 | 19 | 30 | 9 | 8 | 17 | 8 | 0 |
| GLY | 42 | 22 | 8 | 7 | 6 | 8 | 9 | 12 | 1 | 4 |
| HIS | 27 | 25 | 1 | 5 | 4 | 5 | 1 | 3 | 1 | 0 |
| ILE | 69 | 49 | 6 | 7 | 15 | 9 | 10 | 11 | 1 | 2 |
| LEU | 155 | 103 | 14 | 22 | 37 | 14 | 15 | 14 | 10 | 5 |
| LYS | 83 | 52 | 10 | 7 | 26 | 7 | 12 | 10 | 5 | 2 |
| MET | 30 | 13 | 3 | 4 | 7 | 9 | 4 | 7 | 2 | 0 |
| PHE | 48 | 35 | 4 | 3 | 9 | 5 | 3 | 13 | 2 | 1 |
| PRO | 34 | 30 | 2 | 4 | 4 | 5 | 6 | 8 | 3 | 2 |
| SER | 102 | 55 | 15 | 16 | 21 | 9 | 5 | 7 | 2 | 2 |
| THR | 56 | 42 | 6 | 9 | 12 | 7 | 11 | 11 | 3 | 1 |
| TRP | 13 | 8 | 1 | 4 | 4 | 2 | 2 | 1 | 1 | 2 |
| TYR | 32 | 26 | 6 | 11 | 9 | 2 | 3 | 6 | 1 | 0 |
| VAL | 60 | 47 | 4 | 7 | 15 | 6 | 15 | 18 | 3 | 4 |

#### Selection and structural assessment of the AF3 model

To identify a representative AF3 model for subsequent molecular dynamics and metadynamics simulations, the top five AF3 models were first compared with the experimental ATM structure (PDB 7SIC) using PyMOL (Ver 3.1.6.1, student's version from Schdrodinger, LLC) (Supplementary Fig. S1). All five models reproduce the characteristic "butterfly" architecture of the ATM dimer. The FAT–kinase (FATKIN) core shows close structural agreement across the models, whereas the largest differences occur in the Spiral, PRD and FAT-HRD regions, which are also among the regions that are incompletely resolved or structurally flexible in the experimental structure. Thus, the principal differences between the models arise from the completion of structurally less well-defined regions rather than from major differences in the overall ATM architecture.

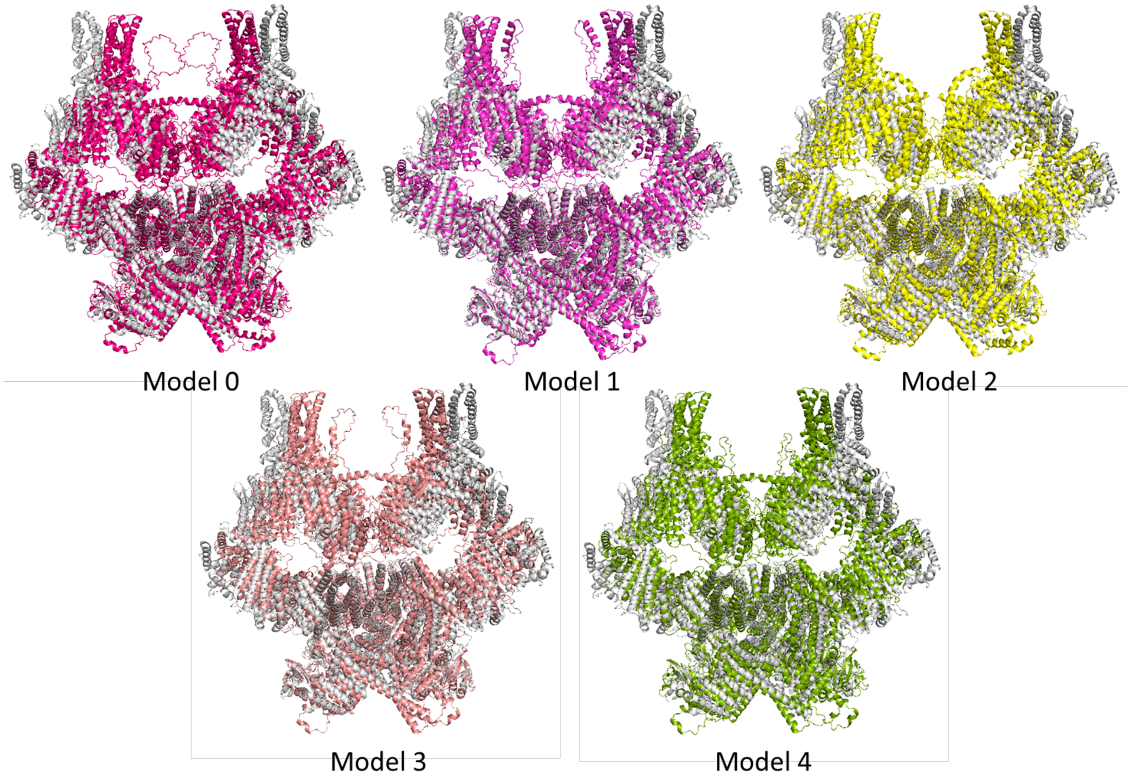


Figure S1: **Alignment of the top five AF3 models of the ATM homodimer with the experimental structure (PDB 7SIC, grey).** Each AF3 model (coloured) is independently superposed on the experimental structure.

Global RMSDs were calculated in PyMOL using the `align` command, which performs sequence-guided structural alignment with iterative outlier exclusion over five refinement cycles. The results are shown in Table S4. Model 1 gives the lowest RMSD at each refinement cycle and the lowest final RMSD (8.33 Å), indicating the closest overall correspondence with the experimental structure among the five models. Domain-wise RMSDs (Table S5) further show that Model 1 has the lowest RMSD for most of the resolved domains, including FAT-TRD1, FAT-TRD2, FAT-TRD3, FAT-

HRD, both kinase lobes, KIN-PRD and FATC. The main exceptions are Spiral and Pincer, which are among the more conformationally variable regions and contain substantial unresolved structure in the experimental model.

Table S4: **Global RMSD (Å) of each AF3 model against the experimental structure (PDB 7SIC), across five iterative PyMOL `align` refinement cycles.**

| | Model 0 | Model 1 | Model 2 | Model 3 | Model 4 |
|---|---|---|---|---|---|
| Cycle 1 rejected atoms | 1193 | 1119 | 1154 | 1508 | 1085 |
| Cycle 1 RMSD | 12.15 | 9.89 | 11.83 | 11.57 | 12.22 |
| Cycle 2 rejected atoms | 572 | 764 | 519 | 1067 | 515 |
| Cycle 2 RMSD | 11.45 | 9.38 | 11.19 | 10.66 | 11.61 |
| Cycle 3 rejected atoms | 303 | 654 | 261 | 781 | 265 |
| Cycle 3 RMSD | 11.19 | 9.09 | 10.96 | 10.17 | 11.37 |
| Cycle 4 rejected atoms | 176 | 650 | 140 | 572 | 152 |
| Cycle 4 RMSD | 11.07 | 8.86 | 10.85 | 9.85 | 11.26 |
| Cycle 5 rejected atoms | 97 | 751 | 70 | 458 | 89 |
| Cycle 5 RMSD | 10.99 | 8.61 | 10.80 | 9.63 | 11.20 |
| **Final RMSD** | **10.955** | **8.329** | **10.769** | **9.456** | **11.164** |

Table S5: **Domain-wise RMSD (Å) of each AF3 model against the experimental structure (PDB 7SIC).**

| Domain | Model 0 | Model 1 | Model 2 | Model 3 | Model 4 |
|---|---|---|---|---|---|
| Spiral | 17.499 | 15.188 | 16.799 | 17.592 | 17.372 |
| Pincer | 9.398 | 7.663 | 8.939 | 8.586 | 9.552 |
| FAT-TRD1 | 5.877 | 4.632 | 6.116 | 5.382 | 6.033 |
| FAT-TRD2 | 2.958 | 2.023 | 3.108 | 2.509 | 3.298 |
| FAT-TRD3 | 5.079 | 2.927 | 5.554 | 4.018 | 5.562 |
| FAT-HRD | 8.471 | 3.706 | 8.955 | 5.457 | 9.137 |
| KIN-Nlobe | 7.865 | 4.364 | 8.718 | 5.906 | 8.465 |
| KIN-Clobe | 7.312 | 2.932 | 7.802 | 3.931 | 7.588 |
| KIN-PRD | 11.669 | 2.642 | 12.087 | 4.044 | 12.020 |
| KIN-FATC | 7.236 | 2.264 | 7.478 | 3.034 | 7.398 |

#### AF3 confidence metrics

The five models were further evaluated using complementary AF3 confidence metrics (Table S6). The *predicted Local Distance Difference Test* (pLDDT) is a per-residue confidence score ranging from 0 to 100 that estimates the accuracy of the local predicted structure, with higher values generally indicating greater confidence. The *predicted Template Modeling score* (pTM) is a global confidence measure ranging from 0 to 1 that estimates the accuracy of the overall predicted structure, including the relative arrangement of domains. The *interface predicted Template Modeling score* (ipTM) is a confidence measure ranging from 0 to 1 for the predicted accuracy of inter-chain interfaces. The AF3 *ranking score* combines pTM and ipTM to rank predicted models according to their overall structural and interface confidence. The *fraction disordered* reports the fraction of residues predicted to be disordered, whereas *Has clash* indicates whether steric clashes are detected in the predicted structure.

Across the five models, the global confidence metrics are closely clustered, with ipTM values of 0.59–0.61, pTM values of 0.60–0.62, and ranking scores of 0.60–0.62. Model 1 falls within these ranges, with an ipTM of 0.59, pTM of 0.60, and ranking score of 0.60, indicating confidence comparable to the other top-ranked models in both overall architecture and predicted inter-protomer interfaces. The predicted fraction of disordered residues is also low (0.02), and no steric clashes are detected. Thus, the AF3 confidence metrics are consistent with Model 1 being a structurally plausible member of the top-ranked solutions.

Table S6: **AF3 confidence metrics for the top five models.**

| Metric | Model 0 | Model 1 | Model 2 | Model 3 | Model 4 |
|---|---|---|---|---|---|
| ipTM | 0.61 | 0.59 | 0.61 | 0.59 | 0.61 |
| pTM | 0.62 | 0.60 | 0.62 | 0.61 | 0.62 |
| Ranking score | 0.62 | 0.60 | 0.62 | 0.61 | 0.62 |
| Fraction disordered | 0.02 | 0.02 | 0.02 | 0.03 | 0.03 |
| Has clash | No | No | No | No | No |

Local confidence was further assessed using average per-domain pLDDT scores (Table S7). FAT-TRD1, FAT-TRD2, FAT-TRD3, KIN-Nlobe, KIN-Clobe and KIN-FATC show consistently higher pLDDT values across the five models (approximately 70–79), whereas Pincer and FAT-HRD show intermediate values ($\sim$69), Spiral shows values of $\sim$66, and KIN-PRD has the lowest values ($\sim$46–49). The same overall pattern is observed across all five models, indicating that local confidence is primarily determined by the structural characteristics of the individual regions rather than by the choice of model. In particular, lower pLDDT values occur in regions containing flexible segments or substantial portions that are unresolved in the experimental structure. This is consistent with the expected uncertainty of local structure in these regions.

Table S7: **Average AF3 pLDDT scores by domain for the top five models.**

| Domain | Model 0 | Model 1 | Model 2 | Model 3 | Model 4 |
|---|---|---|---|---|---|
| Spiral | 65.98 | 65.95 | 65.82 | 65.90 | 65.92 |
| Pincer | 69.08 | 69.19 | 69.21 | 69.18 | 69.20 |
| FAT-TRD1 | 71.04 | 70.58 | 70.98 | 70.97 | 70.81 |
| FAT-TRD2 | 74.57 | 74.69 | 74.69 | 74.82 | 74.71 |
| FAT-TRD3 | 74.38 | 72.44 | 74.45 | 72.56 | 74.40 |
| FAT-HRD | 68.89 | 68.85 | 69.12 | 68.80 | 68.96 |
| KIN-Nlobe | 73.46 | 73.36 | 73.52 | 73.39 | 73.47 |
| KIN-Clobe | 78.07 | 77.10 | 78.10 | 77.10 | 78.07 |
| KIN-PRD | 48.51 | 46.68 | 48.41 | 46.30 | 48.54 |
| KIN-FATC | 78.66 | 75.84 | 78.77 | 75.92 | 78.68 |

#### Domain-wise assessment of the selected model (Model 1)

The combined structural and confidence analysis identified Model 1 as the most suitable representative model for subsequent simulations. Figure S2 shows Model 1 aligned with the experimental structure. The strongest agreement is observed across the FAT–kinase core, while the largest differences occur in regions that are either flexible or incompletely resolved experimentally, including parts of the N-terminal Spiral and C-terminal FAT-TRD3 regions. Model 1 therefore provides close correspondence to the experimentally resolved architecture while supplying a complete structural framework for the regions required for the present mechanistic analysis.

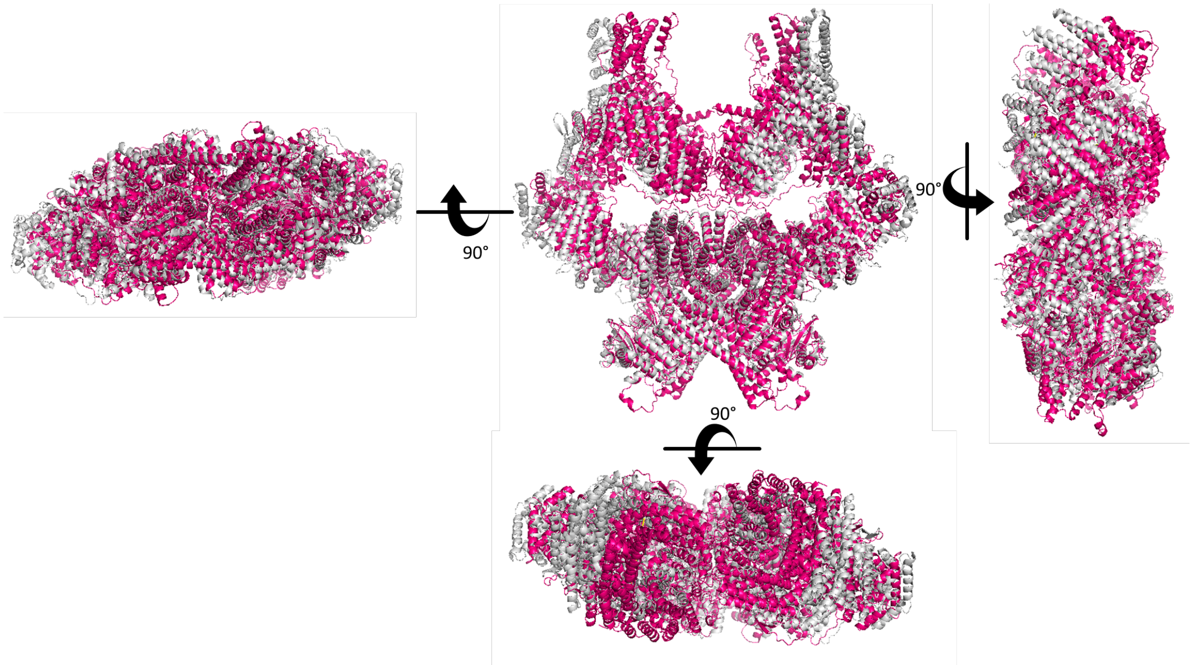


Figure S2: **Multi-angle comparison of AF3 Model 1 (pink) with the experimental structure (PDB 7SIC, grey).**

**Spiral and Pincer.** The Spiral domain (residues 826–877) in Model 1 (blue in Fig. S3) contains a flexible segment with a relatively low pLDDT score of 32.8, compared with 67.2 for the other ordered domains. This is consistent with the conformational variability of this region. The Pincer domain, in contrast, shows good structural agreement with the experimental structure. Model 1 also completes the Spiral–Pincer junction (residues 1166–∼1400) and the region spanning residues 1877–1898, both of which are unresolved in the experimental structure. These reconstructed segments have pLDDT values of approximately 37.6, consistent with structural flexibility in these regions. Importantly, lower pLDDT in such regions does not indicate poor overall model quality; rather, it reflects the absence of a single well-defined local conformation. These are precisely the regions for which AF3 completion provides structural information unavailable from the experimental model.

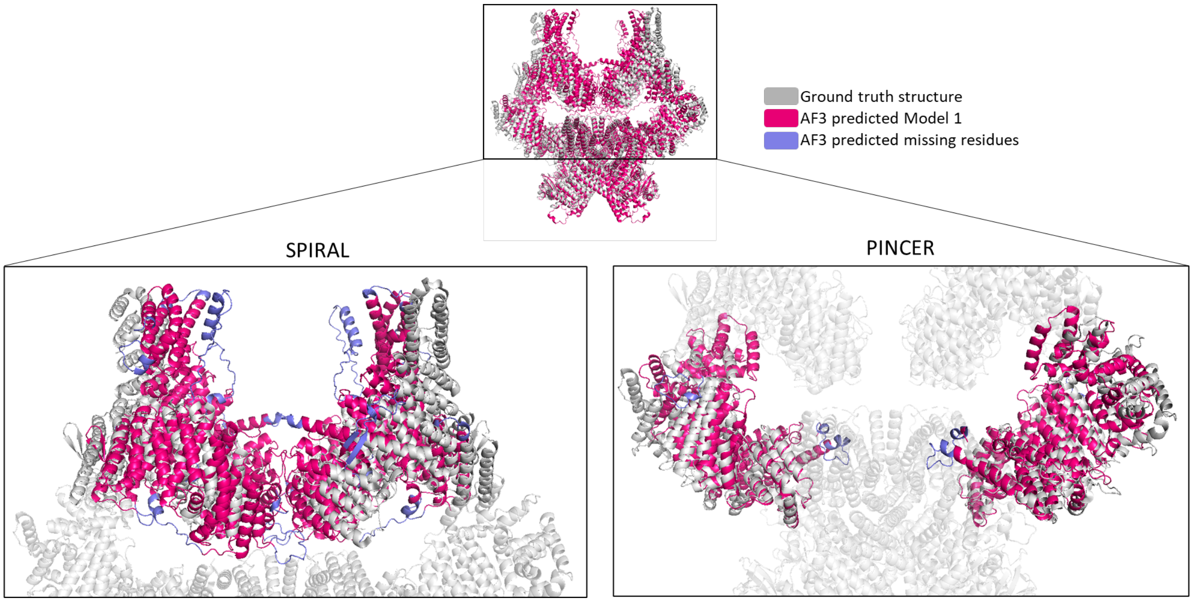


Figure S3: **Spiral and Pincer domains of the experimental structure and AF3 Model 1**, with AF3-predicted missing residues shown in blue.

**FAT-TRD1, TRD2, TRD3 and HRD.** FAT-TRD1, TRD2 and TRD3 show close agreement with the experimental structure (Fig. S4), consistent with the structural stability of these domains and their extensive contribution to the dimer interface. For example, the $\alpha$-helix spanning residues 2074–2088 in each protomer packs closely against its counterpart, with the backbone atoms around Gly2083 separated by only 3.1 Å between protomers, consistent with close inter-protomer packing. Model 1 additionally reconstructs the missing HRD residues, which project away from the main protein body (Fig. S4, blue), providing a plausible surface-exposed region that may contribute to substrate interactions.

**Kinase N-lobe, C-lobe, PRD and FATC.** The kinase domain is highly conserved and fully resolved in the experimental structure. Model 1 reproduces the kinase subdomains with RMSDs of approximately 2–3 Å relative to the experimental structure (Fig. S5), with the slightly larger deviation in the N-lobe localized primarily to loop regions. The $\alpha$-helical elements, N-lobe/C-lobe hinge, and the C-lobe activation and catalytic loops remain well aligned. This agreement is particularly important because these regions form the catalytic core whose structural integrity is central to the subsequent conformational analysis.

Most of the PRD is unresolved experimentally and therefore cannot be directly assessed against a complete experimental structure. PRD has been proposed to occlude the substrate-binding site between the N- and C-lobes, with the FAT-TRD3 domain of the opposing protomer contributing to this occlusion [3, 4]. Model 1 reconstructs both the unresolved PRD and the corresponding FAT-TRD3 region, making this proposed TRD3–PRD interaction directly visible (Fig. S6). The reconstructed interface places the PRD against FAT-TRD3 of the opposing protomer and positions

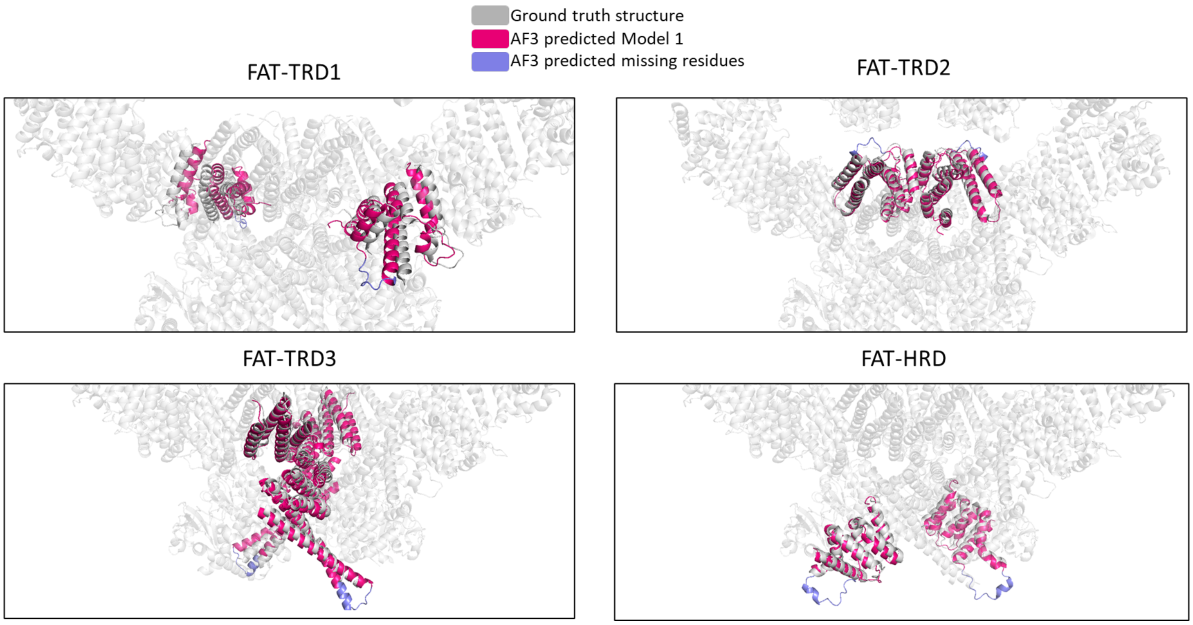


Figure S4: **FAT-TRD1, TRD2, FAT-TRD3 and HRD domains as predicted by AF3 Model 1.**

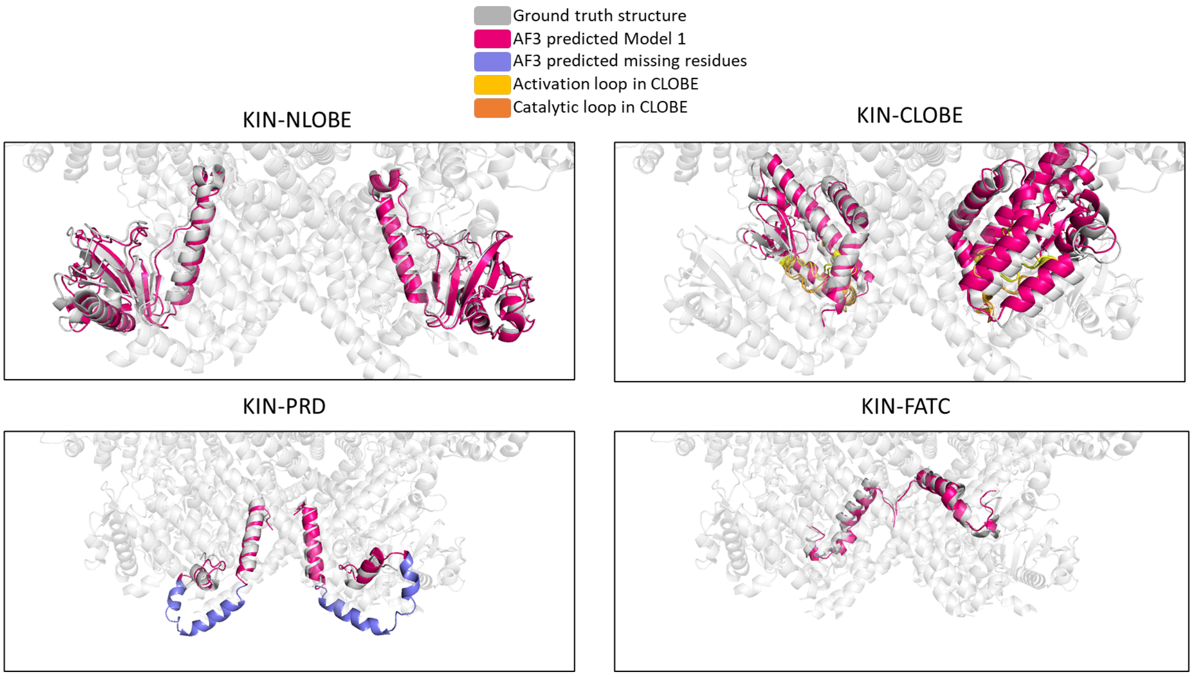


Figure S5: **KIN N-lobe, C-lobe, PRD and FATC domains as predicted by AF3 Model 1.**

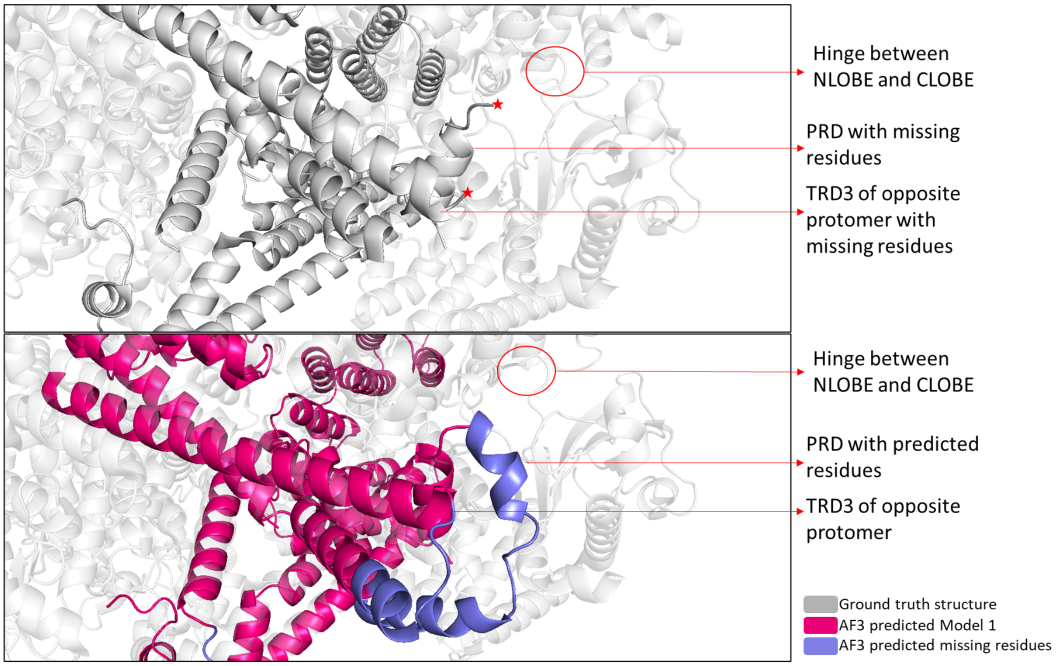


Figure S6: **AF3 Model 1 reconstruction of the FAT-TRD3–PRD substrate-occlusion interaction.** Top: experimental structure (grey), in which both the PRD and FAT-TRD3 of the opposing protomer contain unresolved residues at this interface (red stars). Bottom: AF3 Model 1 (pink), with AF3-predicted residues shown in blue, showing the PRD contacting FAT-TRD3 of the opposing protomer. Together, these domains occlude the substrate-binding groove adjacent to the N-lobe/C-lobe hinge.

both domains across the substrate-binding groove adjacent to the N-lobe/C-lobe hinge. Thus, Model 1 provides a structurally complete framework for examining the proposed substrate-occlusion mechanism, which cannot be resolved directly from the incomplete experimental structure alone.

**Selection of Model 1.** The combined assessment of global and domain-wise structural agreement, AF3 confidence metrics, and domain-level local confidence supports the selection of Model 1 as the representative starting structure. Model 1 has the lowest global RMSD among the five models and shows the closest agreement across most experimentally resolved domains, while its pTM, ipTM and ranking scores remain within the narrow range observed for the other top-ranked predictions. It also contains no predicted steric clashes. Importantly, Model 1 provides a structurally coherent completion of experimentally unresolved regions, including the PRD and FAT-TRD3 interface that is central to the present study. Model 1 was therefore selected as the starting structure for all molecular dynamics and well-tempered metadynamics simulations described in the main text.

#### Supplementary Tables

Table S8: **Domain-wise RMSD comparison of experimental ATM structures (7SIC, inactive [2]; 8OXM, oxidized [3]) and the metastable states F1 (full-length, inactive-like), F6 (full-length, oxidation-compatible) and T1 (Spiral–Pincer-truncated, global minimum).** Structures were superposed on the resolved Cα atoms of FAT-TRD2 from both protomers ("TRD2-anchored"); each domain was then independently superposed and its RMSD recalculated ("independent"). A and B denote the two protomers; T1 lacks Spiral and Pincer domains by construction. All values in Å. This table is referenced in the main text as Supplementary Table S8.

| Reference | Candidate | Domain | TRD2-anch. RMSD (A) | TRD2-anch. RMSD (B) | Indep. RMSD (A) | Indep. RMSD (B) |
|---|---|---|---|---|---|---|
| 7SIC | 8OXM | Spiral | 20.224 | 20.835 | 0.839 | 0.826 |
| 7SIC | 8OXM | Pincer | 10.218 | 10.829 | 1.695 | 1.828 |
| 7SIC | 8OXM | FAT-TRD1 | 6.283 | 6.322 | 1.387 | 1.386 |
| 7SIC | 8OXM | FAT-TRD2 | 0.865 | 0.868 | 0.740 | 0.741 |
| 7SIC | 8OXM | FAT-TRD3 | 3.451 | 3.473 | 2.239 | 2.230 |
| 7SIC | 8OXM | FAT-HRD | 6.825 | 6.887 | 2.018 | 2.022 |
| 7SIC | 8OXM | KIN-Nlobe | 5.836 | 5.898 | 0.917 | 0.916 |
| 7SIC | 8OXM | KIN-Clobe | 7.331 | 7.384 | 0.910 | 0.915 |
| 7SIC | 8OXM | KIN-PRD | 9.773 | 9.859 | 3.496 | 3.587 |
| 7SIC | 8OXM | KIN-FATC | 6.648 | 6.693 | 0.682 | 0.680 |
| 7SIC | F1 | Spiral | 16.313 | 12.233 | 2.354 | 2.023 |
| 7SIC | F1 | Pincer | 9.882 | 5.870 | 1.913 | 2.176 |
| 7SIC | F1 | FAT-TRD1 | 4.042 | 2.288 | 2.017 | 1.579 |
| 7SIC | F1 | FAT-TRD2 | 1.229 | 1.331 | 1.079 | 1.235 |
| 7SIC | F1 | FAT-TRD3 | 2.394 | 3.254 | 1.834 | 2.647 |
| 7SIC | F1 | FAT-HRD | 3.655 | 3.990 | 1.410 | 1.899 |
| 7SIC | F1 | KIN-Nlobe | 3.515 | 4.413 | 1.343 | 1.240 |
| 7SIC | F1 | KIN-Clobe | 3.614 | 1.826 | 1.075 | 1.133 |
| 7SIC | F1 | KIN-PRD | 4.049 | 2.797 | 1.304 | 2.062 |
| 7SIC | F1 | KIN-FATC | 2.970 | 1.734 | 0.630 | 0.809 |
| 8OXM | F1 | Spiral | 7.302 | 12.680 | 2.286 | 2.070 |
| 8OXM | F1 | Pincer | 6.135 | 6.773 | 2.312 | 2.591 |
| 8OXM | F1 | FAT-TRD1 | 3.535 | 5.597 | 1.504 | 1.054 |
| 8OXM | F1 | FAT-TRD2 | 1.374 | 1.296 | 1.289 | 1.234 |
| 8OXM | F1 | FAT-TRD3 | 3.463 | 5.794 | 3.142 | 3.580 |
| 8OXM | F1 | FAT-HRD | 4.824 | 9.010 | 2.244 | 2.454 |
| 8OXM | F1 | KIN-Nlobe | 2.975 | 8.858 | 1.191 | 1.583 |
| 8OXM | F1 | KIN-Clobe | 4.226 | 8.155 | 1.292 | 1.141 |
| 8OXM | F1 | KIN-PRD | 6.836 | 9.860 | 3.727 | 3.971 |
| 8OXM | F1 | KIN-FATC | 3.991 | 7.000 | 1.082 | 0.599 |
| 7SIC | F6 | Spiral | 20.196 | 11.245 | 2.292 | 2.030 |
| 7SIC | F6 | Pincer | 9.090 | 7.835 | 2.590 | 2.785 |

*continued on next page*

| Reference | Candidate | Domain | TRD2-anch. RMSD (A) | TRD2-anch. RMSD (B) | Indep. RMSD (A) | Indep. RMSD (B) |
|---|---|---|---|---|---|---|
| 7SIC | F6 | FAT-TRD1 | 4.621 | 2.753 | 1.781 | 2.079 |
| 7SIC | F6 | FAT-TRD2 | 1.614 | 1.732 | 1.376 | 1.431 |
| 7SIC | F6 | FAT-TRD3 | 3.085 | 3.463 | 2.626 | 2.530 |
| 7SIC | F6 | FAT-HRD | 3.015 | 3.719 | 1.424 | 2.190 |
| 7SIC | F6 | KIN-Nlobe | 3.497 | 5.340 | 1.692 | 1.703 |
| 7SIC | F6 | KIN-Clobe | 4.113 | 1.636 | 1.287 | 1.029 |
| 7SIC | F6 | KIN-PRD | **8.272** | 2.523 | **5.789** | 1.996 |
| 7SIC | F6 | KIN-FATC | 4.017 | 1.505 | 0.772 | 0.657 |
| 8OXM | F6 | Spiral | 6.483 | 18.140 | 2.328 | 2.104 |
| 8OXM | F6 | Pincer | 3.781 | 8.797 | 2.599 | 2.809 |
| 8OXM | F6 | FAT-TRD1 | 2.692 | 6.831 | 1.277 | 1.624 |
| 8OXM | F6 | FAT-TRD2 | 1.788 | 1.729 | 1.554 | 1.527 |
| 8OXM | F6 | FAT-TRD3 | 3.693 | 5.695 | 3.104 | 2.717 |
| 8OXM | F6 | FAT-HRD | 5.862 | 8.631 | 2.488 | 2.566 |
| 8OXM | F6 | KIN-Nlobe | 4.347 | 8.988 | 1.570 | 1.706 |
| 8OXM | F6 | KIN-Clobe | 3.945 | 8.389 | 1.381 | 1.087 |
| 8OXM | F6 | KIN-PRD | 6.929 | 10.449 | 2.852 | 3.743 |
| 8OXM | F6 | KIN-FATC | 3.202 | 7.288 | 1.121 | 1.127 |
| 7SIC | T1 | FAT-TRD1 | 7.690 | 5.129 | 4.417 | 2.650 |
| 7SIC | T1 | FAT-TRD2 | 2.244 | 1.614 | 1.927 | 1.128 |
| 7SIC | T1 | FAT-TRD3 | 3.102 | 3.254 | 2.778 | 2.538 |
| 7SIC | T1 | FAT-HRD | 5.633 | 1.683 | 2.207 | 1.380 |
| 7SIC | T1 | KIN-Nlobe | 5.124 | 2.791 | 2.402 | 1.648 |
| 7SIC | T1 | KIN-Clobe | 3.749 | 2.740 | 1.939 | 1.536 |
| 7SIC | T1 | KIN-PRD | 6.325 | 2.781 | 3.817 | 2.149 |
| 7SIC | T1 | KIN-FATC | 2.844 | 2.361 | 0.881 | 0.959 |
| 8OXM | T1 | FAT-TRD1 | 7.046 | 4.835 | **4.225** | 2.261 |
| 8OXM | T1 | FAT-TRD2 | 2.214 | 1.595 | 1.999 | 1.295 |
| 8OXM | T1 | FAT-TRD3 | 5.403 | 5.551 | **4.386** | 4.180 |
| 8OXM | T1 | FAT-HRD | 8.592 | 7.441 | 2.252 | 2.649 |
| 8OXM | T1 | KIN-Nlobe | 9.198 | 7.024 | 2.723 | 1.992 |
| 8OXM | T1 | KIN-Clobe | 7.477 | 6.763 | 2.087 | 1.483 |
| 8OXM | T1 | KIN-PRD | 10.809 | 9.799 | **4.704** | 4.420 |
| 8OXM | T1 | KIN-FATC | 4.842 | 5.235 | 0.950 | 1.297 |

Table S9: **DBSCAN cluster population per ROI, full-length and Spiral–Pincer-truncated ATM models. “Number of clusters” excludes unclustered noise points (DBSCAN label −1).** Frame counts per cluster are given for clusters 0–5 (blank where a cluster is absent for that ROI); “Total number of frames” is the total number of trajectory frames within the local CV-space neighbourhood of that ROI, including noise points not assigned to any cluster (so cluster frame counts do not necessarily sum to the total). ROI indices correspond to those used throughout the main text and this Supplementary Information (e.g., full-length ROI 1 = F1, ROI 6 = F6; truncated ROI 1 = T1); rows highlighted in the original data (Truncated ROI 1, 4, 6; Full ROI 1) are reproduced here without modification of their significance, which is described where each corresponding metastable state is discussed in the main text.

| Protein | ROI | No. of clusters | Total frames | Cluster 0 | Cluster 1 | Cluster 2 | Cluster 3 | Cluster 4 | Cluster 5 |
|---|---|---|---|---|---|---|---|---|---|
| Truncated | 1 | 4 | 6040 | 718 | 1307 | 3879 | 27 | | |
| Truncated | 2 | 3 | 8030 | 7877 | 56 | 19 | | | |
| Truncated | 3 | 3 | 1223 | 727 | 31 | 122 | | | |
| Truncated | 4 | 3 | 1297 | 183 | 680 | 261 | | | |
| Truncated | 5 | 0 | 364 | | | | | | |
| Truncated | 6 | 1 | 1851 | 1619 | | | | | |
| Truncated | 7 | 1 | 4574 | 4492 | | | | | |
| Truncated | 8 | 1 | 10124 | 10042 | | | | | |
| Truncated | 9 | 3 | 1554 | 129 | 144 | 875 | | | |
| Truncated | 10 | 3 | 22727 | 42 | 22393 | 122 | | | |
| Truncated | 11 | 2 | 7211 | 5248 | 1834 | | | | |
| Truncated | 12 | 6 | 808 | 170 | 124 | 20 | 40 | 28 | 10 |
| Truncated | 13 | 1 | 1863 | 1654 | | | | | |
| Full | 1 | 2 | 15445 | 15301 | 14 | | | | |
| Full | 2 | 3 | 9378 | 9204 | 27 | 22 | | | |
| Full | 3 | 6 | 4983 | 240 | 101 | 28 | 55 | 4199 | 11 |
| Full | 4 | 1 | 10767 | 10763 | | | | | |
| Full | 5 | 1 | 5933 | 5855 | | | | | |
| Full | 6 | 2 | 1330 | 364 | 740 | | | | |
| Full | 7 | 3 | 6752 | 1199 | 39 | 5161 | | | |
| Full | 8 | 4 | 838 | 331 | 39 | 31 | 122 | | |
| Full | 9 | 1 | 8272 | 8204 | | | | | |
| Full | 10 | 3 | 5748 | 1201 | 4339 | 20 | | | |
| Full | 11 | 4 | 842 | 77 | 71 | 102 | 212 | | |
| Full | 12 | 2 | 5397 | 5183 | 53 | | | | |
| Full | 13 | 1 | 1838 | 1597 | | | | | |
| Full | 14 | 4 | 1820 | 193 | 83 | 194 | 1303 | | |
| Full | 15 | 2 | 4620 | 22 | 4495 | | | | |

Table S10: **Domain residue-range definitions used consistently throughout the main text and this Supplementary Information, for a single ATM protomer.** Boundaries follow the convention established for the experimental ATM structure [2] and are shown graphically in main-text Fig. 1. The Spiral–Pincer-truncated construct (T1–T5) retains only the FAT and kinase domains (FAT-TRD1 through KIN-FATC); the Spiral and Pincer domains are absent by construction.

| Domain | Abbreviation | Residue range |
|---|---|---|
| Spiral | Spiral | 1–1166 |
| Pincer | Pincer | 1167–1898 |
| FAT-TRD1 | TRD1 | 1899–2026 |
| FAT-TRD2 | TRD2 | 2027–2193 |
| FAT-TRD3 | TRD3 | 2194–2480 |
| FAT-HRD | HRD | 2481–2614 |
| Kinase N-lobe | Kin-N-lobe | 2615–2771 |
| Kinase C-lobe | Kin-C-lobe | 2772–2958 |
| PRD (PIKK regulatory domain) | PRD | 2959–3027 |
| Kinase FATC | FATC | 3028–3056 |

Table S11: **Summary statistics of residue-level salt-bridge network centrality across the F1, F6 and T1 metastable states.** Degree represents the number of salt-bridge interaction partners, strength represents the sum of edge occupancies for each residue, betweenness measures the extent to which a residue participates in shortest paths through the network, and closeness measures the global accessibility of a residue within its connected network.

| Statistic | Degree | | | Strength | | | Betweenness | | | Closeness | | |
|---|---|---|---|---|---|---|---|---|---|---|---|---|
| | F1 | F6 | T1 | F1 | F6 | T1 | F1 | F6 | T1 | F1 | F6 | T1 |
| Count | 1457 | 1457 | 1457 | 1457 | 1457 | 1457 | 1457 | 1457 | 1457 | 1457 | 1457 | 1457 |
| Mean | 1.6033 | 1.1984 | 0.7014 | 0.6004 | 0.6137 | 0.2573 | 0.000023 | 0.000005 | 0.000358 | 0.000313 | 0.000354 | 0.000246 |
| SD | 0.8875 | 0.7904 | 1.0355 | 0.4979 | 0.5624 | 0.4568 | 0.000070 | 0.000018 | 0.001709 | 0.000275 | 0.000426 | 0.000456 |
| Min | 0 | 0 | 0 | 0 | 0 | 0 | 0 | 0 | 0 | 0 | 0 | 0 |
| 25% | 1 | 1 | 0 | 0.1719 | 0.0500 | 0 | 0 | 0 | 0 | 0.000115 | 0.000042 | 0 |
| Median | 1 | 1 | 1 | 0.5469 | 0.5800 | 0 | 0 | 0 | 0 | 0.000238 | 0.000162 | 0 |
| 75% | 2 | 2 | 1 | 0.9688 | 1.0000 | 0.3807 | 0.000008 | 0.000001 | 0 | 0.000443 | 0.000598 | 0.000336 |
| Max | 6 | 5 | 5 | 2.6875 | 2.9400 | 2.8409 | 0.000866 | 0.000265 | 0.018174 | 0.001650 | 0.002782 | 0.003204 |

Table S12: **State-dependent salt-bridge network centrality of the highest-strength residues in the F1 ensemble.** The 20 residues with the highest strength in the F1 ensemble are shown together with their degree, strength, betweenness and closeness centrality in the F1, F6 and T1 metastable states. Residues are ordered according to their strength in F1.

| Protomer | Residue | Resname | Domain | Degree | | | Strength | | | Betweenness | | | Closeness | | |
|---|---|---|---|---|---|---|---|---|---|---|---|---|---|---|---|
| | | | | F1 | F6 | T1 | F1 | F6 | T1 | F1 | F6 | T1 | F1 | F6 | T1 |
| B | 2895 | GLU | KIN-C-lobe | 3 | 2 | 3 | 2.140625 | 1.99 | 2.840909 | 0.000015 | 0.000003 | 0.002796 | 0.001246 | 0.001864 | 0.001011 |
| B | 2973 | ARG | KIN-PRD | 2 | 1 | 3 | 1.1875 | 0.99 | 2.767045 | 0.000007 | 0 | 0.000106 | 0.000555 | 0.000387 | 0.000958 |
| B | 2642 | ARG | KIN-N-lobe | 1 | 1 | 3 | 0.34375 | 0.03 | 2.75 | 0 | 0 | 0.0002 | 0.00025 | 0.000025 | 0.000382 |
| B | 2016 | ASP | FAT-TRD1 | 4 | 3 | 3 | 2.0625 | 2.09 | 2.551136 | 0.000094 | 0.000023 | 0.000358 | 0.000518 | 0.00024 | 0.000202 |
| A | 3012 | ARG | KIN-PRD | 5 | 5 | 3 | 2.625 | 2.94 | 2.386364 | 0.000075 | 0.00004 | 0.00142 | 0.000592 | 0.002782 | 0.000457 |
| A | 2895 | GLU | KIN-C-lobe | 4 | 3 | 3 | 2.6875 | 2.25 | 2.318182 | 0.000031 | 0.000021 | 0.002796 | 0.000572 | 0.002069 | 0.000458 |
| A | 2444 | GLU | FAT-TRD3 | 2 | 0 | 3 | 0.0625 | 0 | 2.278409 | 0.000007 | 0 | 0.000181 | 0.000108 | 0 | 0.000987 |
| A | 2440 | LYS | FAT-TRD3 | 3 | 2 | 3 | 1.09375 | 1.09 | 2.272727 | 0.000016 | 0.000006 | 0 | 0.00063 | 0.000598 | 0.000951 |
| B | 3012 | ARG | KIN-PRD | 3 | 2 | 3 | 2.109375 | 2 | 2.034091 | 0.000011 | 0.000003 | 0.00004 | 0.001133 | 0.001864 | 0.001006 |
| A | 2832 | ARG | KIN-C-lobe | 2 | 2 | 2 | 2 | 2 | 1.994318 | 0.000001 | 0.000001 | 0.00004 | 0.001452 | 0.001661 | 0.000705 |
| A | 2016 | ASP | FAT-TRD1 | 3 | 2 | 2 | 1.734375 | 1.99 | 1.988636 | 0.00004 | 0.00001 | 0.000075 | 0.00013 | 0.000072 | 0.000611 |
| A | 2929 | ARG | KIN-C-lobe | 1 | 1 | 2 | 0.71875 | 0.99 | 1.988636 | 0 | 0 | 0.000044 | 0.000128 | 0.000071 | 0.000594 |
| B | 2975 | GLU | KIN-PRD | 2 | 1 | 3 | 0.265625 | 0.5 | 1.931818 | 0 | 0 | 0.000252 | 0.000416 | 0.000252 | 0.000918 |
| A | 2175 | ARG | FAT-TRD2 | 3 | 3 | 2 | 1.046875 | 1.11 | 1.931818 | 0.000005 | 0.000021 | 0.000657 | 0.000528 | 0.000657 | 0.000762 |
| B | 2933 | LYS | KIN-C-lobe | 3 | 2 | 3 | 1.953125 | 1.98 | 1.892045 | 0.000028 | 0.00001 | 0.000193 | 0.000485 | 0.000193 | 0.000184 |
| A | 2642 | ARG | KIN-N-lobe | 1 | 0 | 3 | 0.015625 | 0 | 1.829545 | 0 | 0 | 0 | 0.000019 | 0 | 0.000154 |
| B | 2486 | ARG | FAT-HRD | 2 | 3 | 2 | 1.890625 | 2.46 | 1.829545 | 0.000004 | 0.000004 | 0.001791 | 0.000207 | 0.001791 | 0.001972 |
| A | 2336 | GLU | FAT-TRD3 | 2 | 2 | 2 | 2 | 1.92 | 1.829545 | 0.000002 | 0.000003 | 0.000205 | 0.000098 | 0.000205 | 0.003204 |
| A | 1914 | ASP | FAT-TRD1 | 2 | 2 | 2 | 1.5625 | 1.8 | 1.801136 | 0.000005 | 0.000004 | 0.001701 | 0.000653 | 0.001701 | 0.001042 |
| B | 2929 | ARG | KIN-C-lobe | 3 | 2 | 2 | 0.984375 | 1.34 | 1.75 | 0.000089 | 0.00001 | 0.000234 | 0.000525 | 0.000234 | 0.000202 |

Table S13: **State-dependent salt-bridge network centrality of the highest-strength residues in the F6 ensemble.** The 20 residues with the highest strength in the F6 ensemble are shown together with their degree, strength, betweenness and closeness centrality in the F1, F6 and T1 metastable states. Residues are ordered according to their strength in F6.

| Protomer | Residue | Resname | Domain | Degree | | | Strength | | | Betweenness | | | Closeness | | |
|---|---|---|---|---|---|---|---|---|---|---|---|---|---|---|---|
| | | | | F1 | F6 | T1 | F1 | F6 | T1 | F1 | F6 | T1 | F1 | F6 | T1 |
| A | 3012 | ARG | KIN-PRD | 5 | 5 | 3 | 2.625 | 2.940 | 2.386364 | 0.000075 | 0.000040 | 0.001420 | 0.000592 | 0.002782 | 0.000457 |
| A | 860 | ASP | Spiral | 4 | 3 | 0 | 2.078125 | 2.710 | 0 | 0.000152 | 0.000009 | 0.001168 | 0.000315 | 0.000000 | 0 |
| B | 2486 | ARG | FAT-HRD | 2 | 3 | 2 | 1.890625 | 2.460 | 1.829545 | 0.000004 | 0.000004 | 0.001791 | 0.000207 | 0.001791 | 0.001972 |
| A | 2895 | GLU | KIN-C-lobe | 4 | 3 | 3 | 2.6875 | 2.250 | 2.318182 | 0.000031 | 0.000021 | 0.002796 | 0.000572 | 0.002069 | 0.000458 |
| A | 111 | ARG | Spiral | 4 | 3 | 0 | 1.000 | 2.140 | 0 | 0.000126 | 0.000015 | 0.000902 | 0.000300 | 0.000000 | 0 |
| A | 834 | GLU | Spiral | 4 | 3 | 0 | 2.53125 | 2.140 | 0 | 0.000142 | 0.000017 | 0.001320 | 0.000312 | 0.000000 | 0 |
| B | 2016 | ASP | FAT-TRD1 | 4 | 3 | 3 | 2.0625 | 2.090 | 2.551136 | 0.000094 | 0.000023 | 0.000358 | 0.000518 | 0.000240 | 0.000202 |
| B | 110 | ARG | Spiral | 3 | 3 | 0 | 2.250 | 2.060 | 0 | 0.000008 | 0.000011 | 0.001161 | 0.000531 | 0.000000 | 0 |
| B | 1618 | ARG | Pincer | 3 | 3 | 0 | 1.765625 | 2.030 | 0 | 0.000114 | 0.000076 | 0.000507 | 0.000395 | 0.000000 | 0 |
| B | 1489 | ARG | Pincer | 2 | 2 | 0 | 1.28125 | 2.000 | 0 | 0.000135 | 0.000001 | 0.000326 | 0.001661 | 0.000000 | 0 |
| B | 1677 | GLU | Pincer | 2 | 3 | 0 | 1.59375 | 2.000 | 0 | 0.000020 | 0.000032 | 0.000487 | 0.000386 | 0.000000 | 0 |
| B | 1146 | GLU | Spiral | 2 | 2 | 0 | 2.000 | 2.000 | 0 | 0.000003 | 0.000001 | 0.000162 | 0.001661 | 0.000000 | 0 |
| B | 985 | ASP | Spiral | 2 | 2 | 0 | 1.875 | 2.000 | 0 | 0.000008 | 0.000006 | 0.000937 | 0.002207 | 0.000000 | 0 |
| A | 422 | LYS | Spiral | 2 | 2 | 0 | 1.09375 | 2.000 | 0 | 0.000001 | 0.000001 | 0.000249 | 0.001661 | 0.000000 | 0 |
| A | 871 | GLU | Spiral | 2 | 2 | 0 | 1.96875 | 2.000 | 0 | 0.000001 | 0.000001 | 0.001430 | 0.001661 | 0.000000 | 0 |
| A | 106 | LYS | Spiral | 2 | 2 | 0 | 1.8125 | 2.000 | 0 | 0.000000 | 0.000006 | 0.000313 | 0.001239 | 0.000000 | 0 |
| A | 322 | GLU | Spiral | 3 | 2 | 0 | 1.250 | 2.000 | 0 | 0.000004 | 0.000001 | 0.000489 | 0.000371 | 0.000000 | 0 |
| A | 1489 | ARG | Pincer | 3 | 2 | 0 | 1.96875 | 2.000 | 0 | 0.000248 | 0.000022 | 0.000568 | 0.000082 | 0.000000 | 0 |
| B | 2175 | ARG | FAT-TRD2 | 2 | 2 | 2 | 1.703125 | 2.000 | 1.369318 | 0.000030 | 0.000001 | 0.000414 | 0.001661 | 0.000316 | 0.000762 |
| A | 2832 | ARG | KIN-C-lobe | 2 | 2 | 2 | 2.000 | 2.000 | 1.994318 | 0.000001 | 0.000001 | 0.001452 | 0.001661 | 0.000705 | 0.000705 |

Table S14: **State-dependent salt-bridge network centrality of the highest-strength residues in the T1 ensemble.** The 20 residues with the highest strength in the T1 ensemble are shown together with their degree, strength, betweenness and closeness centrality in the F1, F6 and T1 metastable states. Residues are ordered according to their strength in T1.

| Protomer | Residue | Resname | Domain | Degree | | | Strength | | | Betweenness | | | Closeness | | |
|---|---|---|---|---|---|---|---|---|---|---|---|---|---|---|---|
| | | | | F1 | F6 | T1 | F1 | F6 | T1 | F1 | F6 | T1 | F1 | F6 | T1 |
| A | 2895 | GLU | KIN-C-lobe | 4 | 3 | 3 | 2.6875 | 2.25 | 2.318182 | 0.000031 | 0.000021 | 0.002796 | 0.000572 | 0.002069 | 0.000458 |
| A | 3012 | ARG | KIN-PRD | 5 | 5 | 3 | 2.625 | 2.94 | 2.386364 | 0.000075 | 0.000040 | 0.001420 | 0.000592 | 0.002782 | 0.000457 |
| A | 834 | GLU | Spiral | 4 | 3 | 0 | 2.53125 | 2.14 | 0 | 0.000142 | 0.000017 | 0.001320 | 0.000312 | 0 | 0 |
| A | 2973 | ARG | KIN-PRD | 3 | 1 | 3 | 2.4375 | 1.00 | 1.198864 | 0.000014 | 0 | 0.000713 | 0.001258 | 0.000058 | 0.000677 |
| B | 110 | ARG | Spiral | 3 | 3 | 0 | 2.25 | 2.06 | 0 | 0.000008 | 0.000011 | 0.001161 | 0.000531 | 0 | 0 |
| B | 2895 | GLU | KIN-C-lobe | 3 | 2 | 3 | 2.140625 | 1.99 | 2.840909 | 0.000015 | 0.000003 | 0.002796 | 0.001246 | 0.001864 | 0.001011 |
| B | 3012 | ARG | KIN-PRD | 3 | 2 | 3 | 2.109375 | 2.00 | 2.034091 | 0.000011 | 0.000003 | 0.001420 | 0.001133 | 0.001864 | 0.001006 |
| A | 2976 | ASP | KIN-PRD | 3 | 2 | 3 | 2.109375 | 1.95 | 0.903409 | 0.000013 | 0.000007 | 0.000644 | 0.001258 | 0.000058 | 0.000674 |
| A | 860 | ASP | Spiral | 4 | 3 | 0 | 2.078125 | 2.71 | 0 | 0.000152 | 0.000009 | 0.001168 | 0.000315 | 0 | 0 |
| B | 2016 | ASP | FAT-TRD1 | 4 | 3 | 3 | 2.0625 | 2.09 | 2.551136 | 0.000094 | 0.000023 | 0.000331 | 0.000518 | 0.000240 | 0.000202 |
| A | 2933 | LYS | KIN-C-lobe | 3 | 2 | 4 | 2.015625 | 2.00 | 1.715909 | 0.000040 | 0.000017 | 0.000081 | 0.000186 | 0.000121 | 0.000706 |
| B | 2183 | GLU | FAT-TRD2 | 3 | 2 | 1 | 2.015625 | 2.00 | 1.000000 | 0.000079 | 0.000050 | 0 | 0.000500 | 0.000390 | 0.001767 |
| A | 2832 | ARG | KIN-C-lobe | 2 | 2 | 2 | 2.00 | 2.00 | 1.994318 | 0.000001 | 0.000001 | 0.000013 | 0.001452 | 0.001661 | 0.000705 |
| B | 2832 | ARG | KIN-C-lobe | 2 | 2 | 2 | 2.00 | 2.00 | 1.693182 | 0.000001 | 0.000001 | 0.000006 | 0.001452 | 0.001661 | 0.002893 |
| B | 871 | GLU | Spiral | 2 | 2 | 0 | 2.00 | 1.96 | 0 | 0.000001 | 0.000001 | 0.001452 | 0.001627 | 0 | 0 |
| A | 2183 | GLU | FAT-TRD2 | 2 | 2 | 1 | 2.00 | 1.99 | 0.988636 | 0.000153 | 0.000062 | 0 | 0.000356 | 0.000212 | 0.001747 |
| B | 1146 | GLU | Spiral | 2 | 2 | 0 | 2.00 | 2.00 | 0 | 0.000003 | 0.000001 | 0.001661 | 0 | 0 | |
| A | 1677 | GLU | Pincer | 3 | 3 | 0 | 2.00 | 1.46 | 0 | 0.000227 | 0.000035 | 0.000368 | 0 | 0 | |
| A | 2336 | GLU | FAT-TRD3 | 2 | 2 | 2 | 2.00 | 1.92 | 1.829545 | 0.000002 | 0.000003 | 0.000006 | 0.000098 | 0.000205 | 0.003204 |
| B | 850 | GLU | Spiral | 2 | 2 | 0 | 2.00 | 1.52 | 0 | 0.000002 | 0.000064 | 0 | 0.001621 | 0.000240 | 0 |

#### Supplementary Figures

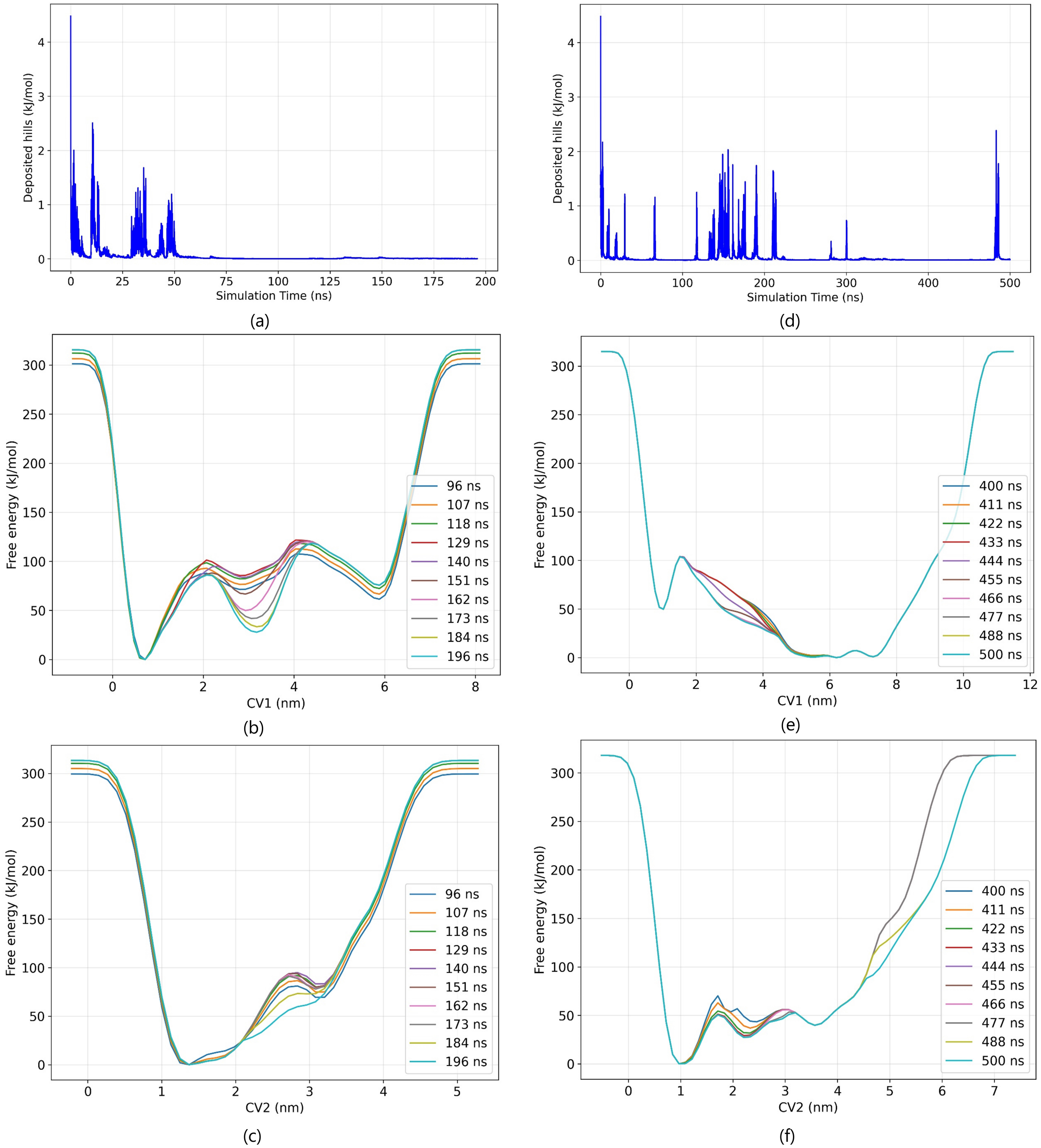


Figure S7: **Stability of the reconstructed free-energy surface over the final 100 ns of each trajectory (a, b, c for truncated protein; d, e, f for full-length protein).** Deposited Gaussian hill height (a, d) and reconstructed one-dimensional free-energy profiles along CV3/CV1 (b, e) and CV9/CV2 (c, f), restricted to the last 100 ns of the truncated and full-length (right) simulations. Profiles from successive time windows within this final segment overlay closely for both systems, indicating that the free-energy surface has stabilized well before the end of each trajectory.

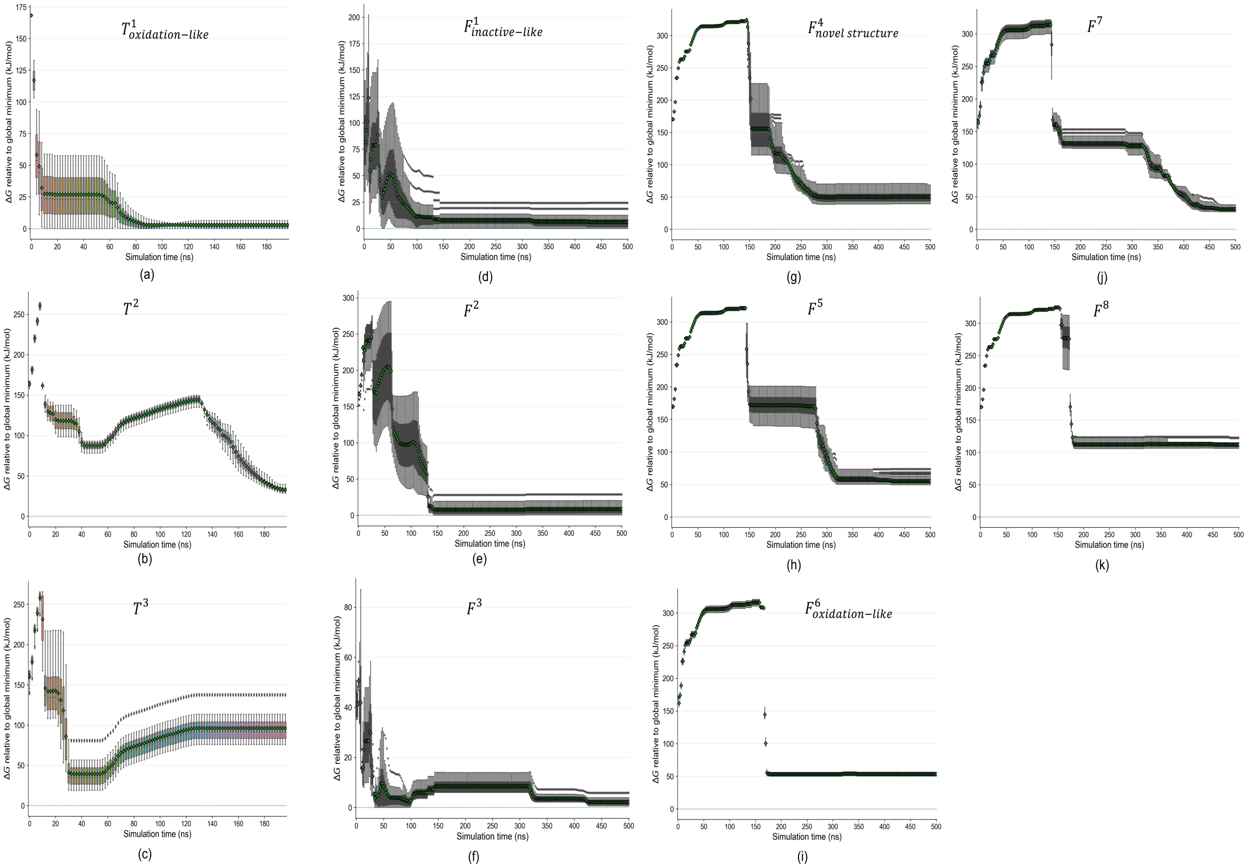


Figure S8: **Convergence of the relative free energies of predefined metastable states.** Free energy of each identified basin relative to the global minimum, plotted as a function of simulation time, for the truncated ATM model states ($T^1_{\text{oxidation-like}}$, $T^2$, $T^3$; panels a–c) and the full-length ATM model states ($F^1_{\text{inactive-like}}$, $F^2$, $F^3$, $F^4_{\text{novel structure}}$, $F^5$, $F^6_{\text{oxidation-like}}$, $F^7$, $F^8$; panels d–k). For each successive FES block, free-energy values were sampled from a local neighbourhood surrounding each predefined minimum (identified by watershed segmentation of the final FES) and summarized as the mean (diamonds) $\pm$ local distribution (boxes and whiskers). Stabilization of these traces to a constant value, together with persistence of the underlying basin, indicates local thermodynamic convergence of the corresponding metastable state.

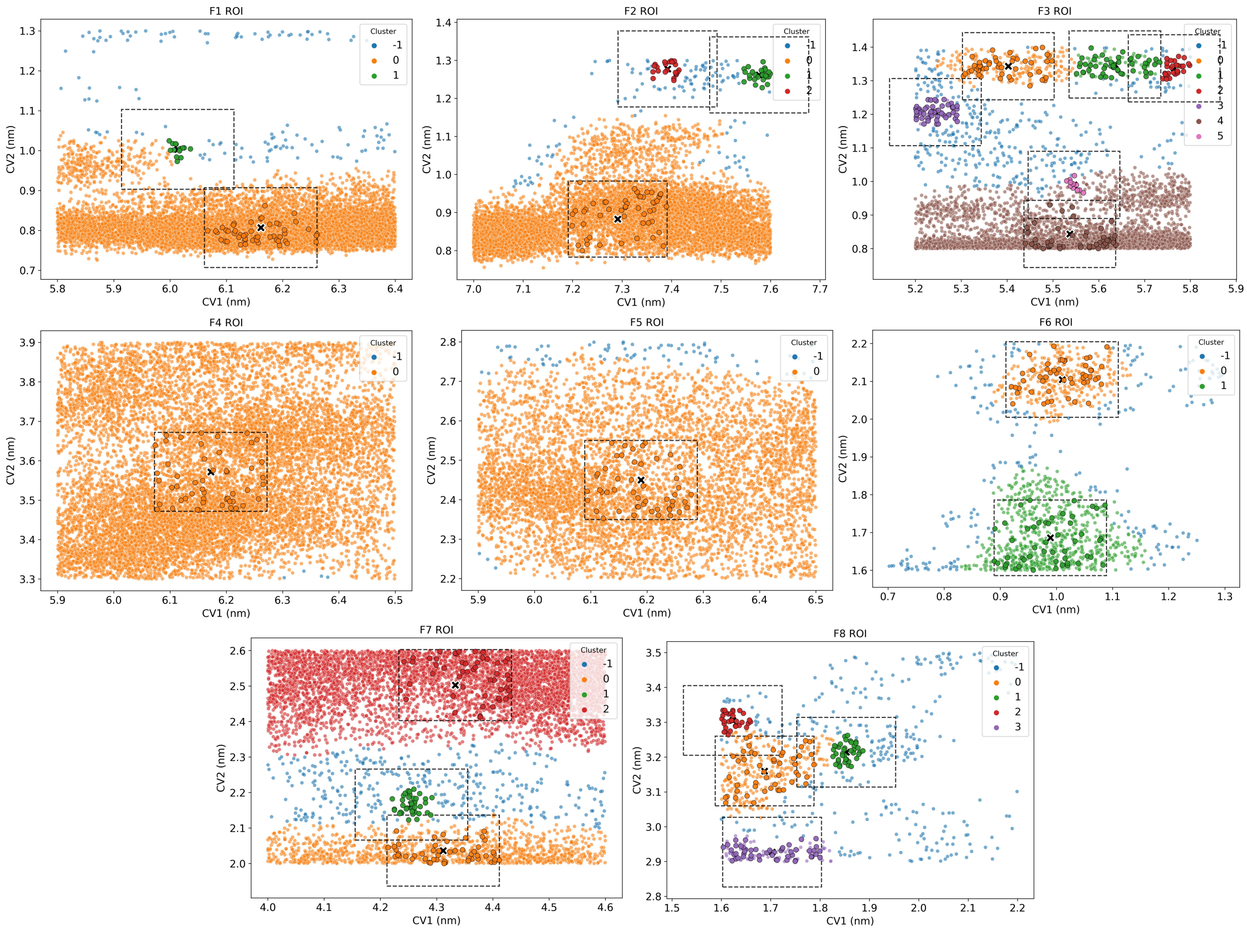


Figure S9: **DBSCAN sub-clustering and representative-structure sampling within each ROI of the full-length ATM model.** Each panel shows trajectory frames within the local CV-space neighbourhood of one ROI (F1–F8), plotted in CV1–CV2 space and coloured by DBSCAN cluster assignment (cluster $-1$, blue, denotes unclustered noise points; clusters $0, 1, 2, \ldots$, other colours, denote resolved sub-populations). Dashed boxes indicate the sampling window defined around each resolved cluster; black crosses mark cluster centroids. Larger, black-outlined points indicate the subset of frames selected as representative structures (50 per ROI, drawn in proportion across all resolved sub-clusters rather than only from the largest sub-population) for the residue-level structural and network analyses described in the main text. ROIs differ substantially in internal population structure: F4 and F5 show a single, broad, continuous population with no resolvable sub-clusters; F6 resolves into two clearly separated sub-clusters; F1–F3, F7, and F8 each contain a principal cluster together with one or more smaller, offset sub-clusters.

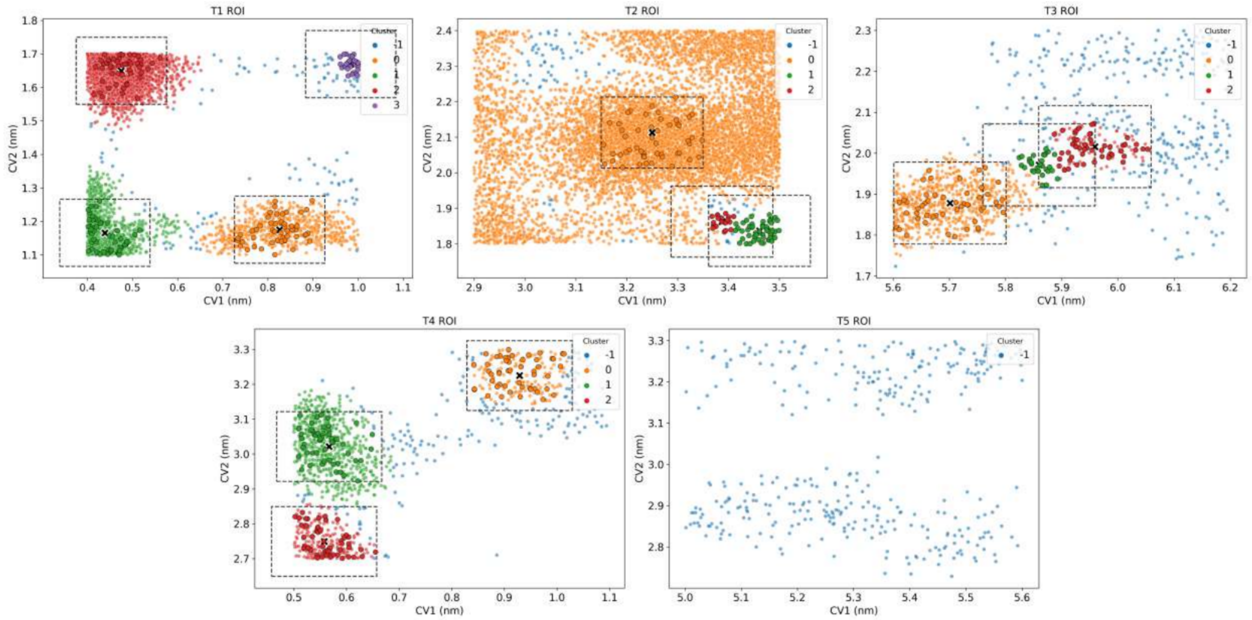


Figure S10: **DBSCAN sub-clustering and representative-structure sampling within each ROI of the Spiral–Pincer-truncated ATM model.** As Supplementary Fig. S10, for the truncated-model ROIs T1–T5. Cluster populations (Supplementary Table S9) mirror the visual structure shown here: T1 resolves into three substantial sub-clusters (718, 1307, and 3879 frames) plus a negligible fourth (27 frames); T2 is dominated by a single large, diffuse population (7877 frames) with two small, tightly co-located minority sub-clusters (56 and 19 frames); T3 resolves into one larger (727 frames) and two smaller (31 and 122 frames) sub-clusters; T4 resolves into three comparably sized sub-clusters (183, 680, and 261 frames); and T5 shows no resolvable sub-cluster structure at all (0 clusters; all 364 frames assigned to noise), consistent with a diffuse, un-clustered population at this ROI.

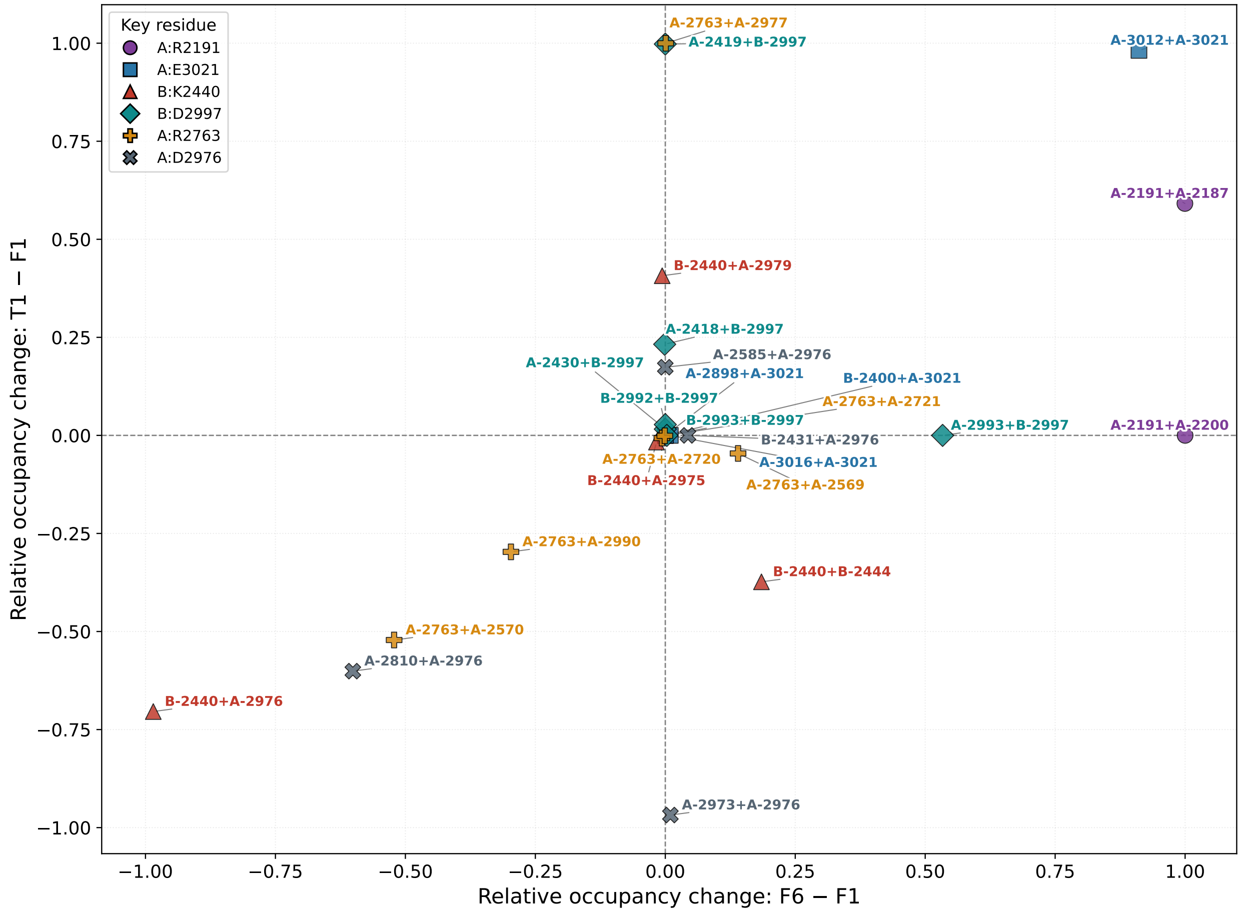


Figure S11: **Pairwise changes in ionic-contact occupancy relative to F1**. Each point represents an ionic-contact residue pair involving the key residues highlighted in Fig. 8 of the main manuscript. The relative change in occupancy in F6 compared with F1 is plotted against the corresponding change in T1 compared with F1. Occupancy denotes the fraction of unbiased-MD frames in which a given contact was observed. The pairwise representation illustrates state-dependent changes in individual ionic contacts and complements the residue-level degree and occupancy analysis by identifying the specific interactions underlying the observed network-level differences.
